# Healing cascades and infections in wounds monitored using a wearable sensor of gaseous flux

**DOI:** 10.64898/2026.06.18.733171

**Authors:** Seunghee Cho, Ansen Tan, Zixi Chen, Kyung Rok Pyun, Shupeng Li, Fujie Yin, Aiwa Zhang, Neena Feldman, Evan J. Neuhart, Amara Devi Moreno, Joyce Eunsung Yoon, Jaeho Shin, Joseph Woojin Song, Jacob Trueb, Yonggang Huang, Guillermo A. Ameer, John A. Rogers

## Abstract

Capabilities for quantitative monitoring of chronic wounds remain an unmet clinical need, as existing diagnostic approaches rely on semiquantitative evaluation of symptoms that lack sensitivity especially during early stages of infection. Here we present a scheme for tracking wound physiology that leverages a miniature, wireless skin-interfaced device for non-contact, transient measurements of the flux of volatile organic compounds (VOCs) and water vapor from the wound microenvironment. Unlike emerging smart bandage platforms that rely on physical contact with the fragile wound bed to interrogate liquid-phase biomarkers, this strategy uses an engineered microclimate and suspended suite of sensors to measure the diffusive transport of wound-derived gases across the wound surface but separated from it. The result enables quantitative evaluation of metabolic activity and healing progression without perturbing the healing tissues. In biofilm growth models of *Staphylococcus aureus*, measurements demonstrate that trends in VOC flux correlate strongly with bacterial growth kinetics and precede any visible biofilm formation. Longitudinal monitoring in infected murine wound healing models shows that concurrent measurements of water vapor and VOC flux provide complementary physiological insights, capturing both the trajectory of barrier restoration and the dynamics of bacterial burden. The findings establish this non-contact sensing scheme as a distinct and clinically translatable paradigm for wound monitoring, with broad implications for non-invasive surveillance of disease states in which tissue metabolic activity and skin barrier integrity serve as actionable physiological readouts.

**Significance Statement:** Limited capabilities in continuous, quantitative assessment of a wound make early diagnosis and effective management challenging, particularly in cases of infection that rapidly progress before symptoms appear. Non-contact approaches for wound monitoring that preserve fragile tissue can transform wound care. In this context, gaseous flux from the wound bed provides an integrative measure of microbial activity and barrier restoration. This study establishes a wearable sensing platform that quantifies these fluxes in real time, enabling early infection detection and temporal tracking of wound healing. These results highlight a path toward personalized treatment strategies and reduced reliance on episodic clinical evaluation.

## Introduction

Chronic wounds remain a major clinical challenge and economic burden (1, 2), partly due to the lack of technologies for identifying subtle signs of emerging complications (1, 3, 4). Early detection of infection is especially critical because bacterial proliferation can progress rapidly to cause persistent inflammation and impede re-epithelialization and extracellular matrix remodeling (5). In medical practice, diagnosis of infection relies largely on visual inspection and assessment of clinical symptoms, including appearance of redness, exudate, and signs of delayed healing (6–8). These signatures, however, only appear after bacteria colonization, at which point the infection has increased resistance to treatment, thereby limiting opportunities for effective intervention. Likewise, microbiological cultures and susceptibility testing provide accurate analysis, but their turnaround times are often long relative to the rapid progression of infections (3, 4, 9). Detection of complications at the earliest stages can significantly improve the outcomes of treatment strategies, which rely largely on antibiotics, despite concerns that their widespread and indiscriminate use may contribute to antimicrobial resistance (10, 11). Accurate evaluation of the efficacy of a selected therapeutic strategy can help prevent antibiotic overuse (11). Episodic clinical follow-ups, often separated by days or weeks, provide only fragmented insights into the wound healing process, with potential delays in important adjustments to care procedures. These challenges underscore the additional need for rapid, frequent and non-invasive monitoring strategies on wound status, especially those that can be used by the patients themselves in the home.

The concept of electronically instrumented wound dressings—so called “smart bandages”—is the subject of considerable research attention as a means of continuous *in situ* wound monitoring. Early demonstrations show that flexible electrochemical sensors integrated into dressing substrates can wirelessly track parameters that shift measurably during infection and inflammation (12, 13), from pH and temperature to uric acid, lactate, hydrogen peroxide, and glucose, as signatures of metabolic activity (14–16). These platforms (12, 14–18), however, require complex engineered systems and direct physical contact with the fragile wound bed and/or with exudate. Disadvantages range from the potential for physical disruptions to healing tissues, to restrictions in wound care approaches, biofouling of the sensors and difficulties in sterilization for cost-effective re-use.

An alternative is in assessments that rely on volatile organic compounds (VOCs), including organic acids, small alcohols, carbonyls, ammonia, and CO2, that diffusively contribute to the microclimate above the wound site (19, 20). Quantitative evaluation of these molecules and the rate of their emergence from the wound can yield important insights into the physiology of the healing tissues for quantitative evaluation of the wound status. Previous studies based on gas sensing demonstrate detection of the concentration of microbial VOCs, but many of these methods involve direct contact of the sensor to the wound tissue and/or require complex instrumentation and time-consuming purging systems (20–23). Means for non-contact, quantitative analysis of gaseous biomarkers *in situ* with a small, convenient device would overcome these challenges. Additional clinical value could follow from simultaneous measurements of the flux of water vapor from the wound, as a basis for tracking the phases of tissue repair that re-establish the structural and functional integrity of regenerated skin at the wound site (1, 24). Because water vapor flux (*fw*) often remains elevated even after clinical closure of the wound relative to that of intact skin, such measurements could also define the integrity of the healed site (25–26).

Here, we report the design and use of a miniature skin-interfaced device capable of non-contact monitoring of molecular gaseous flux for evaluation of wound status and correlation to physiological analysis, without constraining care procedures or occluding diffusive exchange between the wound site with the surrounding ambient. Studies on *in vitro* biofilm growth (168 h), supplemented by rigorous analytical models and microbiological characterization, demonstrate that sharp increases of VOC flux (*fVOC*) correlate strongly with bacterial growth trends and appear prior to any visible biofilm formation. Through long-term monitoring (up to day 30) of infected wounds (*S. aureus*) in mouse models, the data reveal correlations between *fVOC*, bacterial concentration, and biomarkers associated with the inflammatory phase of wound healing, strongly suggesting its potential for early detection of infection, differentiation of infection severity, and evaluation of therapeutic response. Water vapor flux density emerges as a dynamic indicator of skin barrier restoration, capturing progressive changes otherwise quantified through image-based assessments of wound closure and histological analysis.

## Results

The key findings described herein rely on the use of a miniature (Fig. 1*A* and *B* and *SI Appendix* Fig. S1) skin-interfaced device capable of quantifying molecular gaseous flux from wound beds in a non-contacting fashion. As reported in detail elsewhere (27), the device features a chamber that seals against healthy skin around the perimeter of the site of the wound. A valve on the opposite side allows the chamber to diffusively equilibrate with the surroundings when in its open state. In the closed state, water vapor and VOCs from the wound bed accumulate at a rate determined by their characteristic flux. Miniaturized digital sensors within the chamber quantify the time dependence of the corresponding increases and decreases in concentration as the valve closes and opens, respectively, in a cyclic fashion during the monitoring period (Fig. 1*C*). Interactions between a hard magnet and current passing through an electromagnetic coil produce the forces for actuation, while a pair of soft magnets located at the top and bottom of the chamber fix the valve in open and closed states, respectively, to allow for power-efficient, bistable switching. Supporting electronics, including a Bluetooth system-on-chip for wireless communication, enable remote operation powered with a lithium polymer battery (28). The chamber unit includes the suspended sensors, the bistable magnetic valve system, and a disposable adaptor (Fig. 1*D*) that attaches to the bottom of the sensor housing. Disposal of the adaptor after each measurement prevents contamination of the chamber unit through direct contact with the wound area (*SI Appendix* Fig. S2).

**Figure 1.**
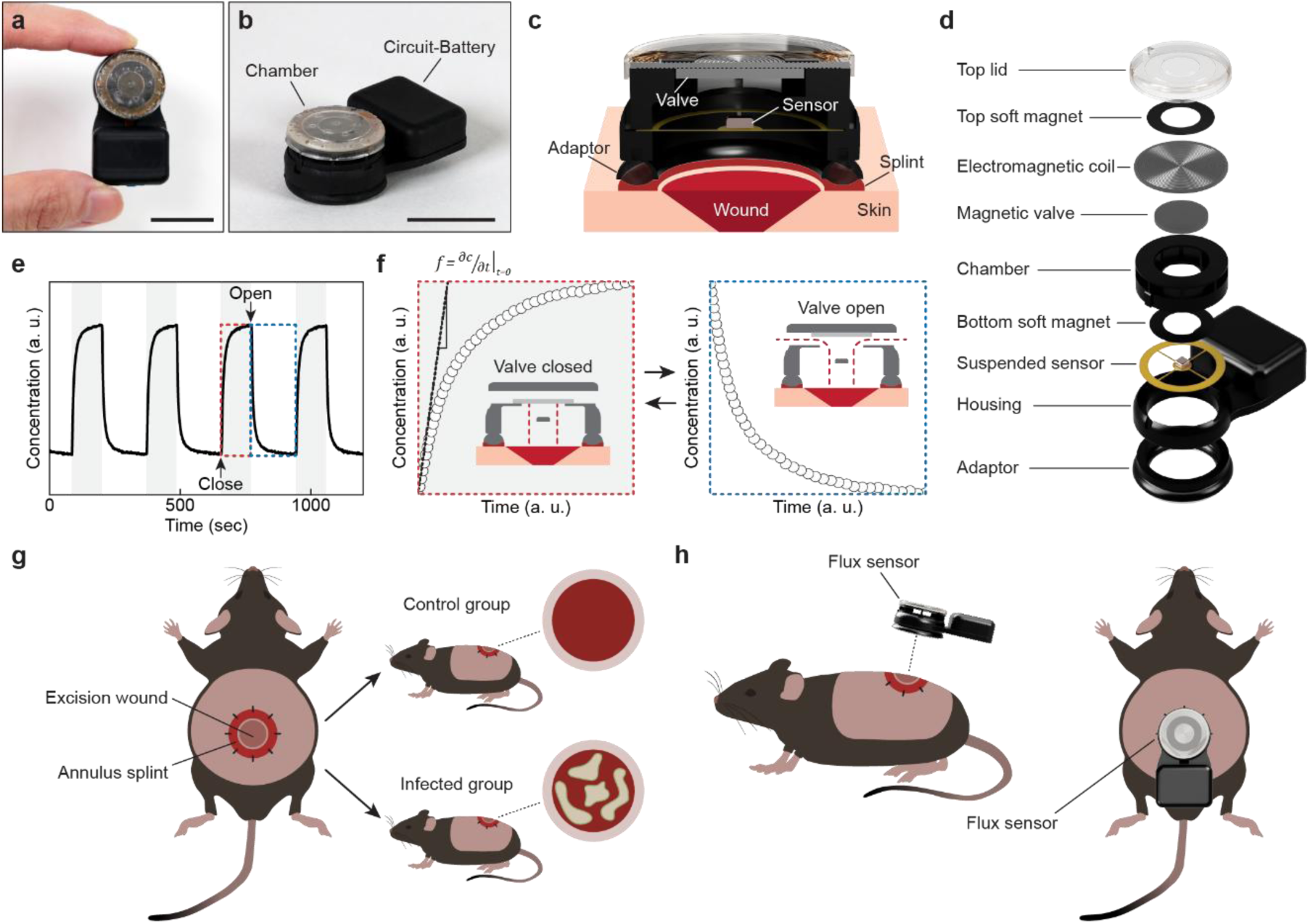
Flux sensor for wound monitoring. (*A–B*), Top view of a flux sensor held between fingertips (*A*) and perspective view featuring the chamber and circuit-battery unit (*B*). Scale bars, 2 cm. (*C*), Cross-sectional view of the analytical chamber of the flux sensor mounted on the wound of the mouse model. (*D*), Exploded view illustration of the sensor featuring the major components. (*E*), Representative data collected through multiple cycles of valve actuation. Shaded regions correspond to periods when the value is closed. (*F*), Accumulation of gaseous species in the chamber while the valve is closed (left) and ventilation while the valve is open (right), corresponding to regions outlined by red and blue dashed lines in e, respectively. The slope of the tangent (black dashed line in graph) to the curve at times immediately after the valve closes corresponds to the flux. Inset images describe the position of the valve and gaseous flow during each phase. (*G*), Use of the flux sensor in mice studies for wound healing monitoring of control and infected wounds. h, Side- and top-view illustrations of the flux sensor placement on the mouse model.

Fig. 1*E* illustrates changes in molecular concentration inside the chamber over time during repeated cycles of valve actuation. When closed, the concentration rapidly increases due to accumulation of molecular vapor released from the wound area and then saturates to an equilibrium value (Fig. 1*F*). Flux corresponds to the change in concentration per unit area per unit time, defined by the slope of the tangent to the concentration plot immediately after the valve closes. After the valve opens, the chamber concentration rapidly decreases to a value defined by an equilibrium with the surrounding environment. Benchtop experiments using water and organic solvents as controlled sources of emission demonstrate that *fw* correlates inversely with the diffusion barrier thickness and that the *fVOC* correlates directly with the source concentration, both quantifiable (*SI Appendix* Fig. S3 and S4). Minimal variation across *fVOC* and *fw* obtained over five independent days also indicates high reproducibility of the valve actuation processes and flux measurements (*SI Appendix* Fig. S5). As considered later in the discussion section, this simple analysis method neglects effects of accumulation of gaseous species within the chamber which can be significant in certain circumstances. Nevertheless, approximate values of flux determined according to the procedures outlined above provide valuable data in tracking changes of interest in the following.

The studies summarized here leverage this technology in *in vitro* and *in vivo* models, the latter with control (non-infected wound) and infected wounds in groups of small animals with single circular excision wounds formed each on their posterior dorsal region to examine the relationship between flux measurements and the wound healing process (Fig. 1*G*). Here, the chamber of the device aligns with the wound area through attachment to an annulus splint held by sutures to the surrounding skin of the wound, providing stability during measurements without contacting or otherwise perturbing the wound bed (Fig. 1*H*).

For the *in vitro* model, the sensor monitors emissions from bacteria cultured on an agar plate. Specifically, the measurements track the behavior of low (103 CFU) and high (107 CFU) concentration bacterial suspensions (*S. aureus*) introduced onto the plates and then cultured in controlled environments, with the sensor fixed on the plate lid aligned with an aperture (Fig. 2*A*). Bacterial load, defined as the total quantity of bacteria present within the biofilm at a given time, exhibits significantly different trends between the two concentrations. 103 CFU shows a continuous increase in bacterial load while 107 CFU shows an initial increase followed by a decrease after 120 h (Fig. 2*B*). Analytical models of bacterial growth kinetics indicate that initial seeding density is a critical parameter because colony geometries can determine nutrient accessibility and spatial constraints. For high inoculation concentrations, closely spaced micro-colonies merge into a quasi-homogeneous film that can be described by Monod kinetics with an added decay term, inducing a later decrease in bacterial load. For low inoculation concentrations, bacteria form large discrete colonies and growth are confined to the colony edge, resulting in the monotonic increase in bacterial load. These models show excellent agreement with the experimentally quantified trends. *fVOC* measurements involve three consecutive cycles of opening and closing the valve, performed once every three hours. The valve otherwise remains in an open state to minimize effects of occlusion. *fVOC* plotted by bacterial load shows a steep increase at low bacterial load and saturate at distinct levels depending on the inoculation concentration (Fig. 2*C*). Analytical models on VOC emission predict a saturation-type relationship due to diffusion-limited release and feedback inhibition by accumulated VOCs and thus verifying the experimental results. Further details on analytical models are described in *SI Appendix*.

**Figure 2.**
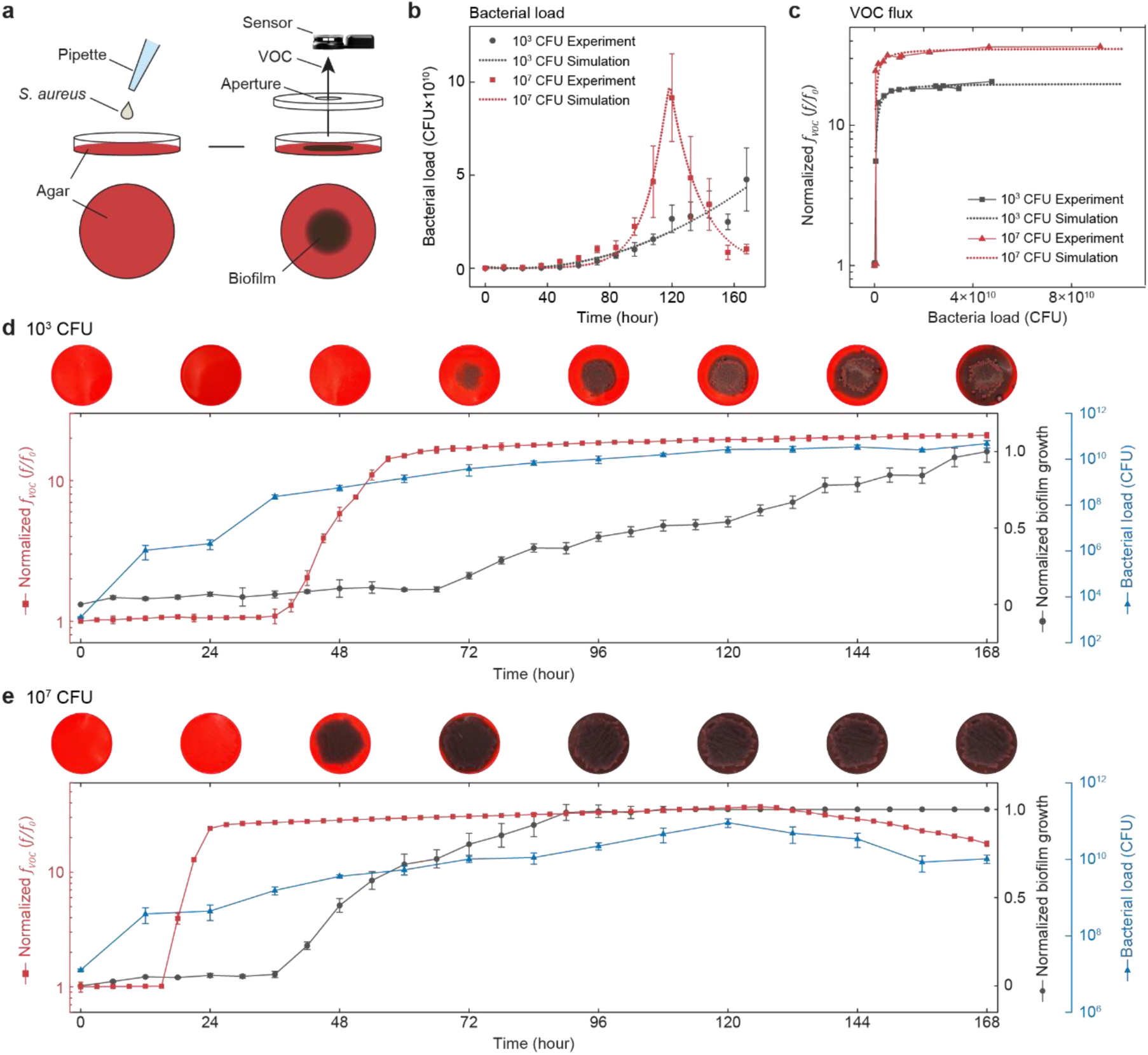
Monitoring VOC flux from an *in vitro* biofilm model. (*A*), Schematic illustration of the experimental set-up for studies of *S. aureus* biofilms grown in Congo red agar plates with the sensor placed on the lid with an aperture. (*B–C*), Experimental (solid line with symbol) and analytical model results (dotted line) for changes in bacterial load as a function of time (*B*) and normalized *fVOC* with increasing bacterial load for inoculation doses (*C*) of 103 CFU (black) and 107 CFU (red). (*D–E*), Images (top) of biofilm plates inoculated with 103 CFU (*D*) and 107 CFU (*E*) taken at 0, 24, 48, 72, 96, 120, 144, 168 h time points and plots (bottom) for normalized *fVOC* (red; *n* = 3), normalized biofilm growth (black; *n* = 4), and bacterial load (blue; *n* = 3) as a function of time. *fVOC* : VOC flux. Data are mean ± s.d.

The use of Congo red agar allows for visualization of the biofilm through color changes from red to black due to reactions with extracellular polymeric substances produced during proliferation of bacteria cells (*SI Appendix* Fig. S6) (29). Analysis of plate images and plots of biofilm growth, bacterial load, and *fVOC* for cultures inoculated with 103 CFU over 7 days show continuous increases in all parameters, although with different onset times (Fig. 2*C*). Increases in bacterial load begin immediately after the start of the experiments. *fVOC* shows a sharp increase beginning at 39 h, roughly 33 h before the biofilm starts to form at 72 h. Cultures inoculated with 107 CFU exhibit similar trends. Specifically, bacterial load increases immediately, followed by a pronounced increase in *fVOC* somewhat later, in this case 18 h, which is 24 h before the biofilm appears at 42 h (Fig. 2*D*). For the 107 CFU inoculation, biofilm entirely fills the plate area at 96 h, leading to accelerated cell death and corresponding decreases in bacterial load and *fVOC* after 120 h and 129 h, respectively. Live/dead stain results reveal that both live cell count and total bacterial load for the 107 CFU inoculation are larger than those of the 103 CFU inoculation at 84 h but smaller at 156 h (*SI Appendix* Fig. S7).

These *in vitro* results suggest the potential for this sensing approach to detect early signs of infection in wounds, as examined here in diabetic mouse models with skin excision wounds (Fig. 3*A*) infected by inoculation with various doses of *S. aureus* (10, 103, and 105 CFU). The experiments involve alignment of the sensor onto the wound (*SI Appendix* Fig. S8) and determination of *fVOC* and *fw* through three consecutive cycles of opening and closing the valve for each measurement session. Additional assessments of the animals include routine determination of wound area, bacteria concentration in the wound, body weight, blood glucose, and wound temperature (*SI Appendix* Fig. S9). These collective measurements occur on healthy skin (HS) before surgery and immediately after surgery (day 0) to establish a baseline and every 3 days thereafter, with subsequent application of wound care including passive cleaning without antimicrobial effects.

**Figure 3.**
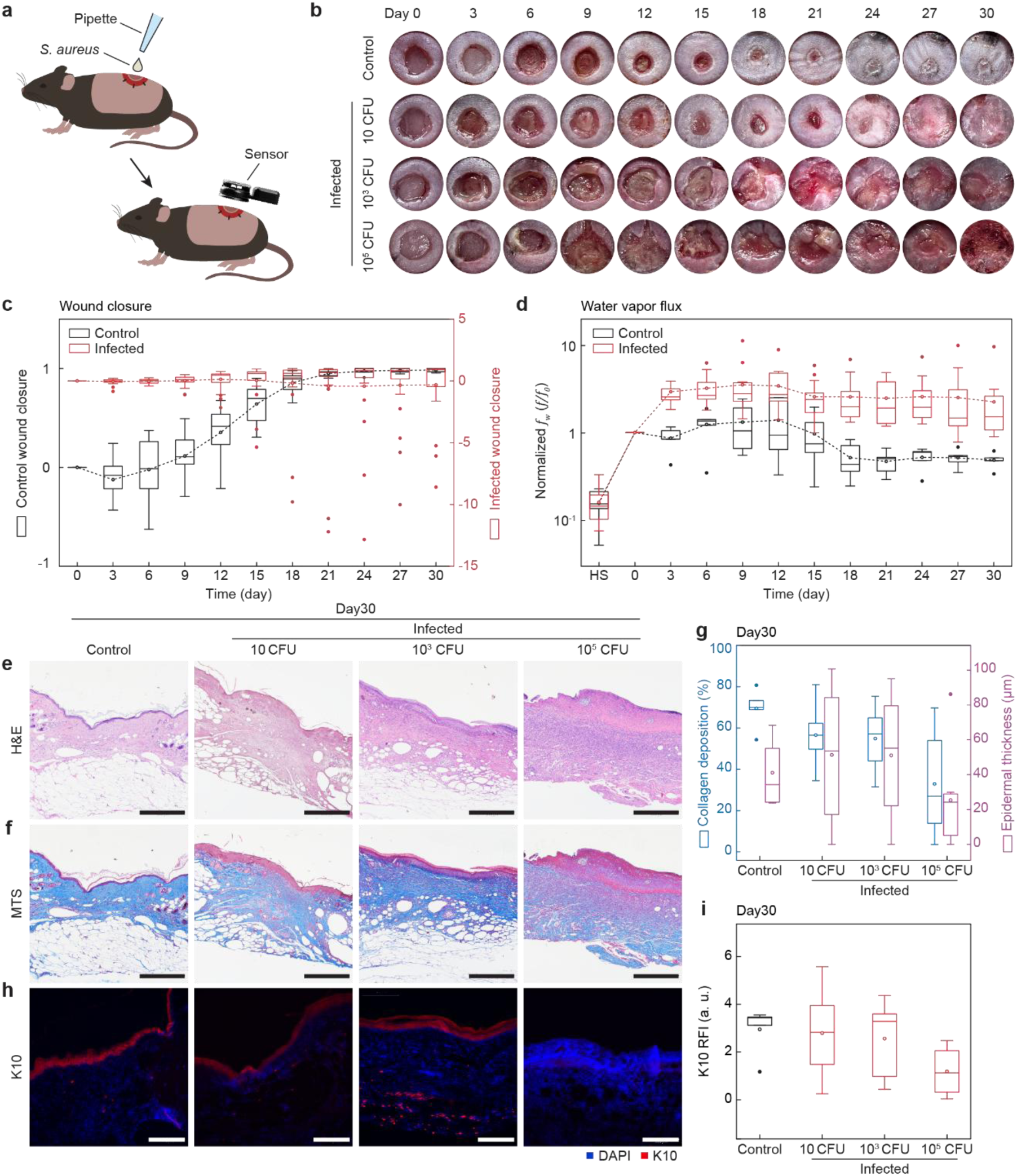
Characteristics of wound healing and re-epithelialization in infected mouse wounds. (*A*), Schematic illustration depicting the infected mouse wound model. (*B*), Representative images of control (non-infected) and infected wounds (bacteria inoculation doses of 10, 103, and 105 CFU) recorded every 3 days. (*C–D*), Wound closure (*C*) and normalized *fw* (*D*) of control (black) and infected (red; 10, 103, and 105 CFU inoculation). *n* = 11 control and *n* = 30 infected for days 3-9, *n* = 8 control and *n* = 27 infected for days 12-18, *n* = 5 control and *n* = 24 for days 21-30 in (*C*). *n* = 5 control and *n* = 15 infected in (*D*). (*E–F*), Representative Haematoxylin and eosin (H&E) staining (*E*) and Masson’s Trichrome staining (MTS) (*F*) images of wound areas on day 30 from control and infected wounds with inoculation doses of 10, 103, and 105 CFU. Scale bars 500 μm. (*G*), Quantitative histological evaluation of collagen deposition (blue) and epidermal thickness (purple) for control (*n* = 5) and infected (*n* = 8 per group) mice. (*H*), Representative immuno-fluorescent staining images for K10 of wound areas on day 30. Scale bars, 200 μm. (*I*), Quantitative histological evaluation of K10 expression on day 30 for control (black; *n* = 5) and infected (red; *n* = 8 per group) mice. For each box plot, the horizontal line within the box represents the sample median and the ends of the box correspond to the interquartile range (first to third quartiles). The whiskers extend beyond the ends of the box by 1.5 × (interquartile range). The open and filled circles represent the mean and outliers, respectively. *fw*: Water vapor flux. HS: Healthy skin. RFI: Relative fluorescence intensity.

Weight loss in control (−18 % by day 30) and infected mice (−29 % by day 30) reflect the degree of systemic distress and metabolic burden (*SI Appendix* Fig. S9*A*). Throughout the 30-day observation period, blood glucose levels remain within 400–600 mg·dL*⁻*¹ (*SI Appendix* Fig. S9*B*) and wound temperatures range from 24–37 °C for both control and infected wounds. Images (*SI Appendix* Fig. S9*C*). Images of the wounds and quantified values for the extent of wound closure highlight differences between the healing processes of control and infected cases (Fig. 3*B* and *C*). On day 3, both control and infected wounds display negative wound closure values (−0.1 for control and −0.05 for infected), indicating an enlargement of the wound, consistent with previous reports on the initial inflammatory phase of wound healing (24, 30, 31). Control wounds approach nearly complete closure by day 21, with an average wound closure of 0.95 and 60 % of all mice resulted in full closure (wound closure value of 1). By contrast, infected wounds exhibit delayed wound closure, with an average value of −0.32 even on day 30. Infected wounds include cases of full wound closure (36 % of all mice) and extreme expansion (average wound closure values of < −5.0). Further comparison between different bacteria inoculation doses reveals that wounds with mild infections (low-inoculation) are more likely to close by day 30 (6 fully closed out of 8 for 10 CFU), whereas severely infected wounds (high inoculation) are more likely to expand (2 expanded out of 8 for 103 CFU and 4 expanded out of 8 for 105 CFU) (*SI Appendix* Fig. S10).

Values of *fw* recorded from wounds reflect the extent of barrier restoration and thus decreases reflect wound closure (Fig. 3*D*) (32, 33). *fw* for control wounds show an increase (days 0-12) to 1.4 times their initial values on day 0 followed by a decrease (days 12-21) and stabilization to half of their initial values. Infected wounds also display similar increases (days 0-12) followed by decreases (days 12-15) and stabilization, but at values that are 2.5 times higher than that on day 0. More specifically, higher inoculation doses show larger increases and stabilization to higher values in *fw*, reaching a maximum of 2.4, 3.2, and 4.6 times their initial values for 10, 103, and 105 CFU, respectively, followed by stabilization to 1.3, 1.6, and 3.8 times their initial values (*SI Appendix* Fig. S11).

Histological analysis of the wound tissue from day 30 reveals delayed re-epithelialization in infected wounds (Fig. 3*E* and *F*). Haematoxylin and eosin (H&E) staining images of mice inoculated with 105 CFU show damaged epidermis layers and excessive formation of scar tissue (Fig. 3*E*), along with decreasing epidermal thickness with increasing inoculation dose for infected wounds (51 μm for 10 CFU, 51 μm for 103 CFU, and 25 μm for 105 CFU) (Fig. 3*G*). Control wounds appear closed with an average epidermal thickness of 41 μm. Masson’s Trichrome staining (MTS) images show that collagen deposition is most pronounced in control mice and significantly reduced in mice with higher inoculation doses (Fig. 3*F*). Quantitatively, collagen deposition on day 30 is greatest for control wounds (70 %) and decreases with increasing inoculation dose in infected wounds (57 % for 10 CFU, 55 % for 103 CFU, and 33 % for 105 CFU) (Fig. 3*G*). Cytokeratin-10 (K10) expression in wounds reflects epidermal maturation and functional restoration of the skin barrier (34). Histological images show notably stronger K10 expression in control mice and almost absent K10 expression in mice inoculated with 105 CFU (Fig. 3*H*). Control wounds show the highest K10 expression (2.9 %), with expression levels decreasing with increasing bacterial inoculation of infected wounds (2.7 % for 10 CFU, 2.6 % for 103 CFU, 1.2 % for 105 CFU) (Fig. 3*I*).

*fVOC* measured in control and infected mice wounds show significantly different trends (Fig. 4*A*). Control wounds exhibit a monotonic decrease in *fVOC*, eventually reaching a value on day 30 that is one tenth of that on day 0. In infected wounds, this parameter rapidly increases during days 0-12, achieving a value on day 12 that is more than fifty times that on day 0, and decreases until day 30 to a value that is twice that of day 0. The bacteria concentration in infected wounds displays a temporal profile similar to that of *fVOC*, with a maximum on day 12 and a decrease until day 30 (Fig. 4*B*, *SI Appendix* Fig. S12). While the temporal profiles vary by mouse due to heterogeneity in host response, the trends of *fVOC* and bacteria concentration show excellent agreement over the 30-day monitoring period for all mice (*SI Appendix* Fig. S13).

**Figure 4.**
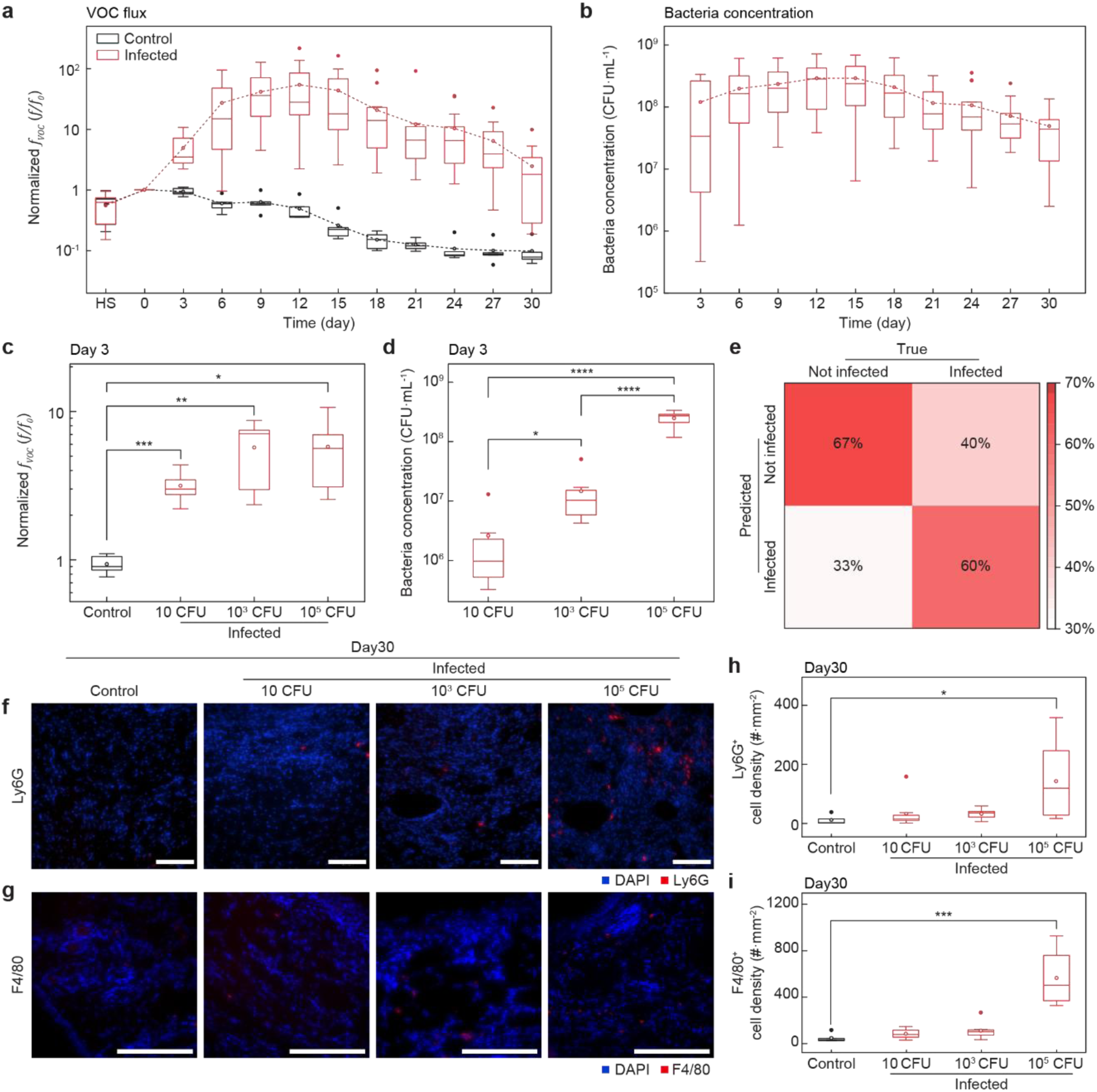
Trends in VOC flux and inflammation in infected mouse wounds. (*A*), Normalized *fVOC* of control (black, *n* = 5) and infected (red, *n* = 15) mice. (*B*), Bacteria concentration of infected mice. *n* = 30 for days 3-6, *n* = 27 for day 9, *n* = 18 for days 12-18, *n* = 15 for days 21-30. (*C*) Normalized *fVOC* from the wound on day 3 for control (black) and infected (red) groups. *n* = 5 per group. (*D*), Bacteria concentration in wounds of mice with inoculation doses of 10, 103 CFU, and 105 CFU on day 3. *n* = 8 for 10 CFU, *n* = 8 for 103 CFU, and *n* = 14 for 105 CFU. (*E*), Confusion matrix summarizing the prediction performance of medical professionals for diagnosing infection in wounds on day 3 through visual assessment only. (*F–G*), Representative immuno-fluorescent staining images for Ly6G (*F*) and F4/80 (*G*) of wound areas on day 30. Scale bars, 100 μm (*F*) and 200 μm (*G*). (*H*), Quantitative histological evaluation of Ly6G+ (red) and F4/80+ (blue) cell density expression for control (*n* = 5) and infected (*n* = 8 per group) mice. For each box plot, the horizontal line within the box represents the sample median and the ends of the box correspond to the interquartile range (first to third quartiles). The whiskers extend beyond the ends of the box by 1.5 × (interquartile range). The open and filled circles represent the mean and outliers, respectively. ****, ***, **, and * indicate *p* < 0.0001, *p* < 0.001, *p* < 0.01, and *p* < 0.05, respectively. *fVOC* : VOC flux. HS: Healthy skin.

Temporal changes in *fVOC* of infected mice with higher inoculation show a prolonged increase and a higher maximum value relative to that on day 0, further supporting that *fVOC* can assist in evaluating the initial infection severity. For inoculation doses of 10 CFU, 103 CFU, and 105 CFU, values continuously increase until days 9, 12, and 15, reaching values 21, 86, and 80 times that of their values on day 0, respectively (*SI Appendix* Fig. S14).

Extremely high bacteria inoculation doses of 107 CFU show outlying trends, where *fVOC* increases reaching at day 12 a value 168 times that on day 0 and then decreases to 19 times the day 0 value, suggesting persistent infection (*SI Appendix* Fig. S15). This behavior is evident through measurements of bacteria concentration, which show elevated values of 1.84×108 CFU·ml-1 on day 30 (*SI Appendix* Fig. S16). The temporal profiles for other parameters, however, resemble those of less severe infections, where wound closure is similar to that of 105 CFU and water vapor flux is similar to that of 10 CFU. This observation can be attributed to the rapid loss of structural integrity in the surrounding skin due to severe infection, which leads to splint detachment and resulted in more advanced wound closure and restoration of skin barrier.

On day 3 since inoculation, infected wounds show significantly different values of *fw* (*p* = 0.00069 for 10 CFU, *p* = 0.0079 for 103 CFU, *p* = 0.051 for 105 CFU) (*SI Appendix* Fig. S17) and *fVOC* (*p* = 0.00031 for 10 CFU, *p* = 0.0058 for 103 CFU, *p* = 0.031 for 105 CFU) from those in control wounds, but the differences for *fVOC* showed stronger statistical significance (Fig. 4*C*). These differences can be associated with the rapid growth in bacteria concentration during the first three days of infection, where inoculations of 10 CFU, 103 CFU, and 105 CFU on day 0 result in 2.6×106 CFU·ml-1, 1.5×107 CFU·ml-1, and 2.5×108 CFU·ml-1, respectively, on day 3 (Fig. 4*D*). Evaluation of wound images through visual inspection on day 0 and 3 by medical professionals yielded an accurate assessment in only 64% instances (Fig. 4*E*). Differences in bacterial growth rate by inoculation dose are most prominent during early stages of infection, but these differences reduce over time, with bacteria concentrations converging to the same order of magnitude by day 30, where inoculations of 10 CFU, 103 CFU, and 105 CFU on day 0 result in 3.0×107 CFU·ml-1, 5.0×107 CFU·ml-1, and 6.7×107 CFU·ml-1, respectively (*SI Appendix* Fig. S18). This finding suggests that even initially mild infections can intensify to levels comparable to more severely infected wounds, without appropriate intervention.

Immunofluorescence analysis of tissue on day 30 allows for further investigation of the effects of bacterial infection on the wound healing status (Fig. 4*F* and *G*). Ly6G and F4/80 are each markers for neutrophils and mature macrophages which are predominant during early inflammatory phases and active immune response phases of wound healing, respectively (35, 36). Staining images show that both Ly6G and F4/80 expressions are negligible for control wounds and increase with increasing inoculation dose for infected wounds. Quantitatively, infected wounds with high inoculation (105 CFU) show significantly higher Ly6G expressions compared to those of controls (*p* = 0.048) (Fig. 4*H*). Similarly, F4/80 expressions for the 105 CFU inoculation case are substantially higher than those of the control (*p* = 0.00050) (Fig. 4*I*). Ly6G and F4/80 expressions for infected wounds with low bacteria inoculation doses (10 and 103 CFU) are comparable to those of control (*p* > 0.05). Blood analysis results also indicate prolonged inflammation in animals with bacterial infections as per higher total white blood cells (3.1×104 μl-1) and increased proportions of neutrophils (51 %) compared to those of the control (1.7×104 μl-1 white blood cells and 23 % neutrophils) (*SI Appendix* Fig. S19).

Further studies that compare infected wounds with an inoculation dose of 105 CFU (standard wound care group) to those with the same inoculation dose treated with hydrochlorous acid (antimicrobial wound care group) demonstrate the potential utility of *fVOC* in the assessment of inflammation (Fig. 5*A*). *fVOC* in the standard wound care group shows a continuous increase over days 0-15 reaching a maximum of 80 times that on day 0 and decreases to a value that is 1.5 times that on day 0. In contrast, the antimicrobial wound care group exhibits a rapid increase in *fVOC* with a higher maximum on day 6 (87 times that on day 0) followed by a continuous decrease to a lower value that is 0.41 times that on day 0. Bacteria concentration also differs between the two groups. The standard wound care group displays a maximum on day 15 followed by an overall decrease, while the antimicrobial wound care group shows a monotonic decrease until day 30 (Fig. 5*B*).

**Figure 5.**
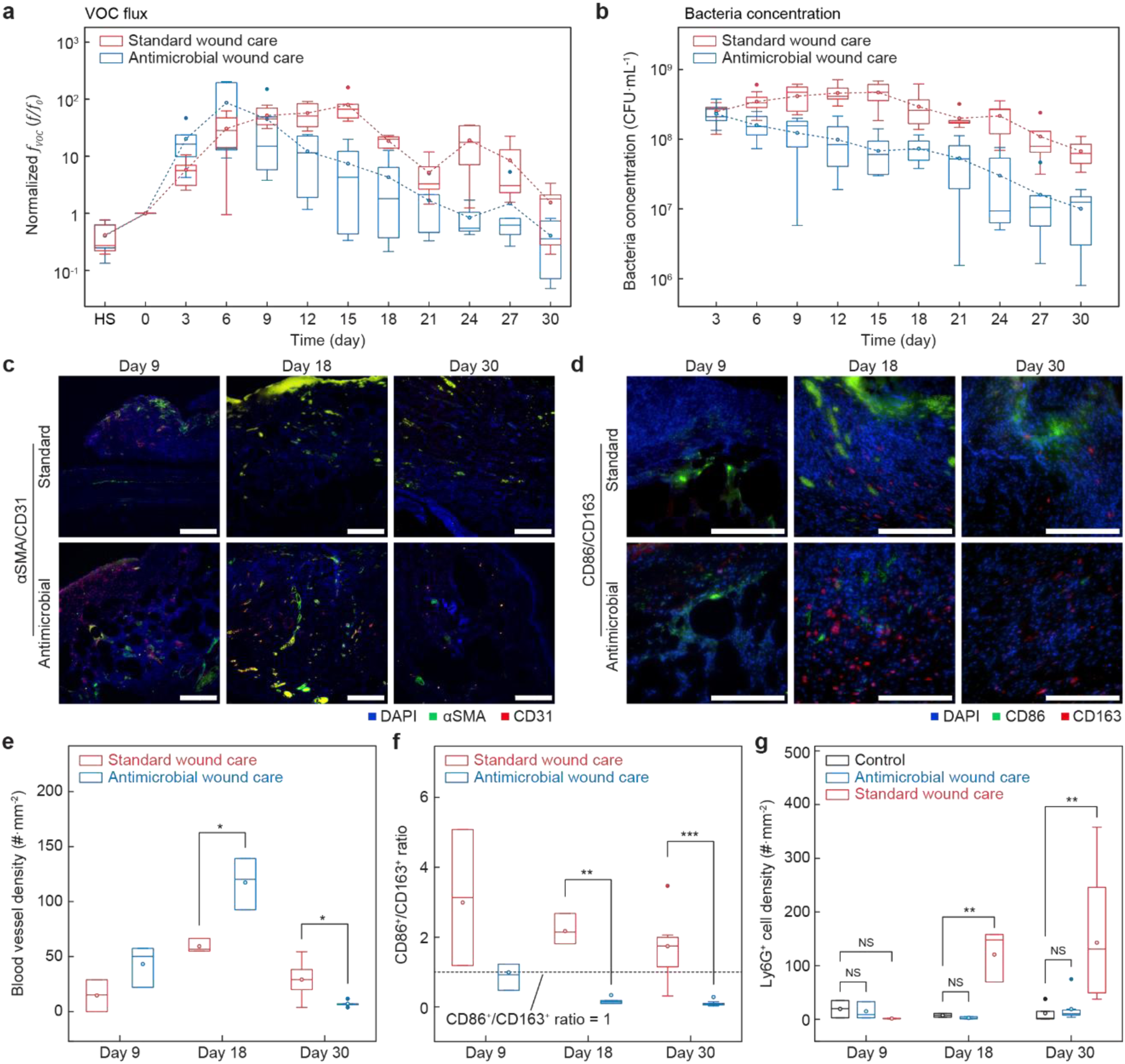
Influence of antimicrobial treatment of infected wounds on healing and VOC flux. (*A–B*), Normalized *fVOC* (*A*), and bacteria concentration (*B*) from infected wounds with inoculation doses of 105 CFU with standard wound care (black; *n* = 5) and antimicrobial treatment (blue; *n* = 5). (*C–D*), Representative immuno-fluorescent staining images for αSMA/CD31 (*C*) and CD86/CD163 (*D*) of wound areas at time points of day 9, 18, and 30. Scale bars, 200 μm. **e-f**, Quantitative histological evaluation of blood vessel density (*E*), CD86+/CD163+ ratio (*F*), and Ly6G+ cell density (*G*) on days 9, 18, and 30. *n* = 3 control and n = 3 infected mice per wound care group for days 9 and 18. *n* = 5 control and *n* = 8 infected mice per wound care group for day 30. For each box plot, the horizontal line within the box represents the sample median and the ends of the box correspond to the interquartile range (first to third quartiles). The whiskers extend beyond the ends of the box by 1.5 × (interquartile range). The open and filled circles represent the mean and outliers, respectively. ***, **, *, and NS indicate *p* < 0.001, *p* < 0.01, *p* < 0.05, and p > 0.05, respectively. *fVOC* : VOC flux. HS: Healthy skin.

Antimicrobial treatment of the wounds significantly enhances wound healing as per quantification of blood vessel and macrophage markers via immunofluorescence (Fig. 5*C* and *D*). Cells positive for αSMA and CD31 visualize blood vessel density at the wound site (37, 38). Qualitatively, both standard and antimicrobial wound care groups show strongest αSMA/CD31 expressions on day 18 compared to days 9 and 30 (Fig. 5*C*). Quantitatively, blood vessel densities of the antimicrobial wound care group on day 18 and 30 are significantly higher (*p* = 0.015) and lower (*p* = 0.033), respectively, than those of the standard wound care group (Fig. 5*E*). Additionally, CD86 and CD163 each stain for polarized phenotypes of macrophages that are pro-inflammatory and pro-regenerative, respectively (39, 40). Staining images exhibit higher expressions of CD86 in the standard wound care group and increased expressions of CD163 in the antimicrobial wound care group across days 9, 18, and 30 (Fig. 5*D*). Analysis of the CD86+/CD163+ cell ratio reveals significant differences between the two groups (*p* = 0.0012 for day 18 and *p* = 0.00042 for day 30), where macrophages present in the standard wound care group are predominantly pro-inflammatory (CD86+/CD163+ > 1) while those in the antimicrobial wound care group are primarily pro-regenerative (CD86+/CD163+ < 1) (Fig. 5*F*). Temporal analysis of Ly6G+ cells indicates that the antimicrobial wound care group exhibits expression levels comparable to those of control wounds, whereas the standard wound care group shows significantly higher expressions (*p* = 0.0053 for day 18 and *p* = 0.0090 for day 30) (Fig. 5*G*). Blood analysis further supports these findings, with control and antimicrobial wound care groups maintaining similar levels below 2.0 ×104 μl-1 across days 9, 18, and 30, while the standard wound care group displays a continuous increase up to 4.0×104 μl-1 (*SI Appendix* Fig. 20). Consistent trends are observed in wound closure and *fw* measurements, where the antimicrobial wound care group closely resembles control wounds. (*SI Appendix* Fig. S10, S11, and S21).

## Discussion

The results presented here demonstrate that continuous monitoring of gaseous flux from wound beds yields data with strong relationships to epidermal barrier integrity and biological processes that govern infection and wound healing progression. An advanced version of a non-contact, wireless wearable platform for quantitating gaseous flux yields measurements of *fVOC* that indicate a divergence between behaviors of infected and non-infected wounds at early time-points. These findings indicate that molecular emissions from wounds contain clinically relevant information before the appearance of more prominent clinical signs of infection (22, 41). As complementary information, temporal variations in *fw* provide insight into the recovery of skin barrier function during wound healing. Skin excision disrupts the stratum corneum and increases water loss through loss of lipid barrier integrity (32, 42). As wound closure progresses, keratinocyte migration and proliferation drive re-epithelialization, while subsequent maturation of keratinocytes restores the epidermal structure (33, 34). Decreasing *fw* therefore reflects maturation of the regenerated epidermis and restoration of barrier properties. Infected wounds, however, exhibit persistently elevated flux, consistent with delayed epithelialization and altered epidermal architecture associated with sustained inflammatory stress (43, 44). Histological features further support these functional observations, including differences in epidermal thickness and delayed epithelial maturation in more severely infected wounds (K10). Together, these findings suggest that molecular flux – in the form of both *fVOC* and *fw* -- from the wound bed represent a previously underexplored class of physiological biomarkers that integrate microbial metabolism, immune activity and structural recovery within the wound microenvironment (27, 45–47).

The mechanistic basis of these flux signatures arises from interactions between bacterial metabolism, immune response and structural remodeling of the skin barrier. VOCs emerge during early stages of bacterial proliferation as metabolic by-products generated during active growth and enable detection of microbial activity before macroscopic biofilm formation (48, 49). As infection progresses, host immune responses further influence the chemistry of the wound.

Recruitment of innate immune cells introduces oxidative and inflammatory pathways that generate additional VOCs through lipid peroxidation and reactive oxygen species–mediated reactions, reflecting sustained inflammatory activity within the wound bed (Ly6G, F4/80) (50, 51). In parallel, *fw* reflects the structural progression of epidermal repair, as mentioned above.

Persistent infection disrupts these processes through inflammatory signaling and bacterial toxins that impair keratinocyte proliferation and matrix remodeling (43, 44). Elevated *fw* in these cases therefore reflects prolonged exposure of hydrated dermal tissue and incomplete maturation of the stratum corneum. Simultaneous monitoring of *fVOC* and *fw* thus provides powerful insights into the physiological status of the wound, capturing both microbial activity and the structural restoration of tissue.

The same principles allow evaluation of treatment efficacy. Suppression of bacterial proliferation through antimicrobial intervention reduces microbial metabolism and leads to rapid decreases in *fVOC*. Concurrently, histological analysis of immunofluorescent markers indicates a transition toward a regenerative wound environment, accompanied by normalization of inflammatory markers (Ly6G expression and total white blood cell counts) and convergence of *fw* and wound closure toward control levels. Resolution of inflammation accompanies improved vascularization and tissue remodeling, processes that enhance oxygen and nutrient delivery to the regenerating tissue and facilitate progression toward the remodeling phase of healing (αSMA, CD31) (37, 38). Because transitions between inflammatory and regenerative phases are critical indicators of whether wounds progress toward healing or chronicity, the correspondence between these biological processes and flux measurements serve a physiologically meaningful role in wound status assessment during therapy.

The enabling technology is uniquely well suited for this application because it largely decouples measurements of flux from absolute concentrations, thereby significantly improving the robustness against baseline drift and environmental variability. Under high flux conditions, however, the flux values determined from the tangential slope may not fully account for progressive accumulation of gas molecules within the chamber when ventilation is insufficient for complete diffusive equilibration. This phenomenon can shift the baseline (concentration during valve-open) over successive cycles, causing deviations between measured and true, unobstructed molecular flux. Although limiting the number of measurement cycles (*in vivo*) and extending ventilation times (*in vitro*) can mitigate these effects, further studies incorporating full transient analysis of concentration–time profiles may improve the quantitative accuracy of flux values determined from the recorded data. The strong temporal correspondence between *fVOC* and bacterial load together suggests that infection-related processes dominate the measured signals under the conditions explored here.

Future introduction of chemical selectivity through integration of high-dimensional sensor arrays with multivariate models may enable deconvolution of overlapping transport phenomena, enabling differentiation of bacterial species and polymicrobial compositions based on characteristic emission patterns. Addition of collection reservoirs can extend the capabilities of this platform further for *ex vivo* analysis (e.g., GC–MS), linking real-time flux measurements with molecular identification. Translation of this sensing technology to clinical settings may involve integration into wound dressings or wearable bandage platforms (46, 47, 52). Wireless transmission of data to clinicians from use in outpatient settings may reduce the need for frequent clinical visits (16, 53). Further studies in large animal models and clinical trials will be necessary to determine how this technology and associated measurements translate to human wounds, which exhibit greater variability in size, microbiome composition, and physiology.

## Materials and Methods

### Flux sensor fabrication

The sensors consist of a flexible printed circuit board (f-PCB), surface mount electronic components, magnetic elements for the valve system, a Li-polymer battery, and structural frame. The electronic components bonded to the f-PCB include a Bluetooth low energy system on chip (BLE SoC, ISP1807, Insight SIP, Sophia Antipolis, France), a sensor that measures humidity and VOC (BME680, Bosch, Reutlingen, Germany), and a planar electromagnetic coil. The magnetic elements include a neodymium magnet (diameter 11.3 mm; height 0.9 mm) as the valve and two soft magnets. The soft magnets use 20 wt% magnetite nanoparticles (50-100 nm, Merck, Burlington, MA, USA) mixed with polydimethylsiloxane (PDMS; Sylgard 184, Dow, Midland, MI, USA; 10:1 mixing ratio) and spin-cast at 400 rpm, thermally cured at 50 °C for 12 h and laser cut to define annulus shapes (inner diameter 8.4 mm; outer diameter 14.2 mm). 3D printing (Form3+, Formlabs, Somerville, MA, USA) defined the structural components including the chamber and encapsulation. For the disposable adaptors, polydimethylsiloxane (PDMS; Sylgard 184, Dow; 10:1 mixing ratio) was poured into molds defined by 3D printing, degassed, and cured. The adaptors were sterilized by spraying 70% isopropyl alcohol and air dried before use. Attachment to the bottom side of the sensor chamber unit was achieved through an interference fit.

### Flux calculation

Flux, defined as the change in concentration per unit area per unit time, is calculated based on the slope of the tangent to the concentration immediately after valve closing as

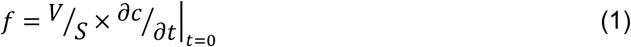

where *V* is the chamber volume, *S* is the area of the analytical region, *c* is the molecular concentration of the chamber, and *t* is time.

### Biofilm model

Agar plates were prepared with 10 ml of brain heart infusion agar (Merck, Burlington, MA, USA), which consists of agar (15.0 g/L), brain extract (7.8 g/L), dextrose (2.0 g/L), disodium phosphate (2.5 g/L), heart extract (9.7 g/L), proteose peptone (10.0 g/L), sodium chloride (5.0 g/L), Congo red dye (0.8g), and deionized water (1L). *S. aureus* (ATCC 23235, Manassas, VA, USA) was thawed overnight in Luria-Bertani (LB) broth at 37 °C and 225 rpm. Inoculation involved introduction of 10 μl suspensions containing approximately 103 and 107 CFU of *S. aureus* (*SI Appendix* Table S1). Plates were kept upside down, in the dark, at temperatures of 18-23 °C. Plate images were taken every 6h using a custom light box setup and a digital SLR camera (Canon EOS 6D, Oita, Japan). For quantification of the bacterial load of biofilm, the entire agar and biofilm was removed from the plate, dissolved in phosphate-buffered saline, and vortexed gently for 1-2 min to allow for homogenization. Concentrations were calculated using a Grant bio Den-1B densitometer and a LIVE/DEADTM *Bac*LightTM kit (Thermo Fisher Scientific, Waltham, MA, USA) was used to fluorescently stain for bacterial viability. Plates for imaging and bacteria quantification were distinguished batches from those for VOC measurement to minimize influence from intervention.

### Mice

Male diabetic mice (BKS.Cg-m +/+Leprdb, 00642; homozygous for Leprdb) aged 8-12 weeks were obtained from Jackson Laboratory (Bar Harbor, ME, USA). The Institutional Animal Care and Use Committee at Northwestern University’s Center for Comparative Medicine approved all animals and procedures before experimentation (IS00018748).

### Infected wound model

To anesthetize the mice, a portable gas induction and inhalation system (RC2 Roden Circuit Controller, VetEquip, Livermore, CA, USA) delivered 2.5% isoflurane in oxygen. The fur on the dorsal side was then removed. Once asleep, injections of Meloxicam (20 mg/kg) and Bupivacaine (2 mg/kg) were administered subcutaneously, followed by addition of 6-0 nylon sutures (Ethicon, Raritan, NJ, USA) to attach an annulus acrylate splint (inner diameter, 10 mm; outer diameter, 12 mm) to prevent skin contraction. A full-sized dermal excision was then created to define the wound using a 6-mm diameter punch biopsy (Acuderm, Fort Lauderdale, FL, USA). For the infected wounds, a 10 μl suspension of *S. aureus* (10, 103, 105, and 107 CFU) was added to the wound bed (*SI Appendix* Table S2). Preparation of *S. aureus* inoculums were prepared as described previously. Occlusive dressings (TegaDerm, 3M, Maplewood, MN, USA) were used to cover the wound area.

### Post-operative care

Mice were housed individually with access to standard laboratory chow and water ad libitum. The animal facility maintained a 14 h:10 h light/dark cycle, ambient temperatures of 18-23 °C, and humidity levels between 40 and 60%. Approximately 24 h following initial wound creation, a second dose of Meloxicam (20 mg/kg) was delivered subcutaneously. Wound care was performed every 3 days utilizing a dry swab to remove and collect exudate, followed by cleansing using a gauze pad soaked with hypochlorous acid (Vashe, Urgo Medical North America, Fort Worth, TX, USA) for the antimicrobial wound care (AWC) group and with saline for all other infected wounds (SWC: standard wound care). Wound area, weight loss, blood glucose concentration, and wound temperature were monitored prior to wound care. Wound area was characterized by three blinded observers using ImageJ (1.54p). Blood glucose concentrations were measured from whole venous blood using a blood glucose monitor (Contour next, Ascensia, Basel, Switzerland) with a measurement upper limit of 600 mg·dL-1. Wound temperatures were determined using a handheld infrared camera (Flir Edge Pro, FLIR, Goleta, CA, USA) and the FLIR Thermal Studio software (v.2.0.58).

### Blood tests

Once euthanized, blood samples were collected by making an incision at the nape of the neck and stored in EDTA-coated tubes to prevent coagulation. Samples were transferred to the Northwestern University Center for Comparative Medicine for complete blood count (CBC) analysis with leukocyte differential. For hematological analysis, targeted traits included hemoglobin concentration (HGB), hematocrit (HCT), mean corpuscular volume (MCV), mean corpuscular hemoglobin (MCH), mean corpuscular hemoglobin concentration (MCHC), red blood cell (RBC) count, RBC distribution width (RDW), and total white blood cell (WBC) count. The concentration of WBC subtypes (neutrophils, lymphocytes, monocytes, eosinophils, and basophils) were also measured.

### Bacteria quantification

Before each flux measurement, wounds were gently swabbed with sterile dry cotton tipped applicators (Puritan®, Guilford, ME, USA) three times in a circular motion ensuring coverage of all sides of the applicator with wound exudate. The collected swab samples were then stored briefly in 1 mL PBS solution. Immediately following wound care, swab samples were vortexed at 3200 rpm for 30 s (Analog Vortex Mixer, FisherbrandTM, Waltham, MA, USA) to allow for thorough homogenization of the wound exudate. Serial dilutions of the exudate solution were performed at a 1:10 ratio in PBS. Dilutions were cultured onto nutrient agar (DifcoTM, BD, Franklin Lakes, NJ, USA) using disposable inoculating loops (FisherbrandTM, Waltham, MA, USA) and incubated overnight at 37 °C. Agar plates containing a countable range (30-300 CFUs) of colonies were used for quantifying bacterial concentration using the following equation:

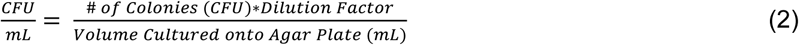

### Tissue processing and sampling

The mice were euthanized by carbon dioxide inhalation followed by cervical dislocation. The skin tissue was excised and fixed with 4 % paraformaldehyde for 24 h. The fixed tissues were then washed with PBS for 48 h and then gradually dehydrated to be embedded in paraffin. The embedded tissues were sectioned into slides with 5 μm thickness for histological analysis. Brightfield histology, including H&E (Sigma-Aldrich, Burlington, MA, USA; GHS332 and HT110116) and Masson’s Trichrome (Abcam, Cambridge, UK; ab150686) staining, was performed to assess epidermal thickness, granulation tissue formation, and collagen deposition. Immunofluorescence staining was conducted to evaluate vasculature using rabbit anti-CD31 antibody (Abcam, Cambridge, UK; ab182981) and mouse anti-αSMA antibody (Abcam, Cambridge, UK; ab7817), keratinocyte differentiation using mouse anti-cytokeratin-10 antibody (Invitrogen, Waltham, MA, USA; MA5-13705), macrophage presence using rat anti-F4/80 antibody (Invitrogen, Waltham, MA, USA; 14-4801-82), macrophage polarization using rat anti-CD86 antibody (Invitrogen, Waltham, MA, USA; 14-0862-82) and rabbit anti-CD163 antibody (Abcam, Cambridge, UK; ab182422), and neutrophil presence using rat anti-Ly6G antibody (BD, Franklin Lakes, NJ, USA; 551459). ImageJ (version 2.16.0/1.54p) was used for quantitative analysis.

### Histological analysis

Images of wound tissues were acquired using an inverted fluorescence microscope (Eclipse Ti2-E, Nikon, Tokyo, Japan) equipped with NIS-Elements software. For each tissue, two fields were imaged at 10× for H&E staining, one field at 10× for Masson’s trichrome staining, three fields at 20× for CD31/α-SMA and K10 immunofluorescence staining, and four fields at 40× for CD86/CD163, F4/80, and Ly6G immunofluorescence staining. Epidermal thickness was quantified from H&E-stained images by averaging measurements taken at six locations distributed along the wound site, with an average spacing of approximately 500 µm. Collagen deposition was quantified from Masson’s trichrome–stained images by color deconvolution of the blue channel and measurement of the non-background area fraction using a fixed threshold. Blood vessel density was quantified by manually counting CD31*⁺*/α-SMA*⁺* lumen-containing structures normalized to the area of granulation tissue in each image. Cytokeratin-10 fluorescence was quantified by measuring the mean gray value of the Cytokeratin-10 channel within the epidermal layer and subtracting the mean gray value measured from a negative control region. CD86/CD163, F4/80, and Ly6G positive cell densities were quantified by manually counting DAPI-positive nuclei that exhibited overlapping fluorescence signals for the corresponding marker, followed by normalization to tissue area. ImageJ (version 2.16.0/1.54p) was used for all the quantitative histological analysis.

### Clinical survey for visual inspection of wound infection

Medical professionals (*n* = 8) were recruited to complete a single-blinded survey administered *via* an online link. The survey consisted of 15 questions, each presenting a pair of wound images from mouse models, including one image obtained immediately after surgery (day 0) and a corresponding image from day 3. For each pair, participants were asked to determine whether the wound on day 3 appeared infected. The dataset included a mixture of control and infected wounds, and participants were not informed of the proportion or identity of infected cases, ensuring blinding to ground truth labels. Responses were aggregated and evaluated using a confusion matrix to quantify accuracy of classification.

### Data processing

Calculation of flux from VOC concentration and humidity data involved regression fitting based on the exponential form of

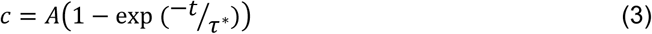

where *c* is the VOC concentration or humidity, *t* is time, and *A* and *τ\** are variables.

For normalized flux in *in vitro* and *in vivo* studies, each value (*f*) was divided by its baseline value (*f0*) which is from 0 h immediately after inoculation and day 0 immediately after surgery, for *in vitro* and *in vivo* studies, respectively, and therefore presented as normalized quantities (*f/f0*) to express the relative change with respect to the baseline value.

Biofilm growth was quantified as Euclidean distance in RGB space calculated as

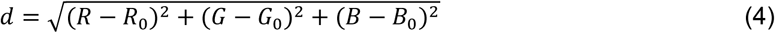

where (*R*, *G*, *B*) and (*R0*, *G0*, *B0*) are the average plate color at each time point and at 0 h, respectively. Wound closure corresponds to the percentage healed compared to day 0 calculated as

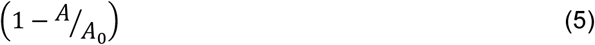

>where *A* and *A0* are the wound area at each time point and on day 0, respectively.

### Statistical analysis

One-way ANOVA tests (SPSS) were used to derive two-sided *p* values. Minimum sample sizes for experiments were not predetermined.

## Acknowledgments

The authors thank Prof. Hartman for providing laboratory space for in vitro experiments; Dr. Steve Xu for useful discussions. This work is supported by the Querrey-Simpson Institute for Bioelectronics, the Center for Advanced Regenerative Engineering (CARE) at Northwestern University, Ameer Wound Healing: R01DK131302, and the CZ Biohub Spoke Project – Agreement 2/4/2025.

## Author Contributions

S.C., A.T., G.A., and J.A.R. conceived the project. S.C. designed the device hardware; fabricated devices (with assistance from A.Z., N.F., and J.E.Y), performed benchtop and in vitro testing/characterization, and all flux data analysis (with assistance from J.S. and J.W.S.). A.T. performed in vitro testing/characterization (with assistance from A.Z.), designed the surgical procedure, conducted surgeries, post-operative care, routine measurements, histology sample preparation, blood collection, and blood analysis (with assistance from A.D.M.). Z.C. conducted analysis of qualitative and quantitative histology (with assistance from N.F. and E.J.N.). K.R.P. fabricated devices and performed benchtop testing/characterization. S.L. and F.Y. developed analytical models. J.T. developed firmware. S.C., A.T., G.A., and J.A.R. wrote the paper. All authors read and provided comments on the paper. Y.H., G.A., and J.A.R. jointly supervised the work.

## References

1. V. Falanga, et al., Chronic wounds. Nature Reviews Disease Primers 8, 1–21 (2022).

2. X. Deng, M. Gould, M. A. Ali, A review of current advancements for wound healing: Biomaterial applications and medical devices. Journal of Biomedical Materials Research Part B: Applied Biomaterials 110 (2022).

3. M. J. Carter, et al., Chronic wound prevalence and the associated cost of treatment in medicare beneficiaries: changes between 2014 and 2019. Journal of Medical Economics 26, 894–901 (2023).

4. C. K. Sen, et al., Human skin wounds: a major and snowballing threat to public health and the economy. Wound repair and regeneration : official publication of the Wound Healing Society [and] the European Tissue Repair Society 17, 763–71 (2009).

5. S. L. Percival, S. M. McCarty, B. Lipsky, Biofilms and Wounds: An Overview of the Evidence. Advances in Wound Care 4, 373–381 (2015).

6. R. G. Frykberg, J. Banks, Challenges in the Treatment of Chronic Wounds. Advances in Wound Care 4, 560–582 (2015).

7. M. Reddy, S. S. Gill, W. Wu, S. R. Kalkar, P. A. Rochon, Does This Patient Have an Infection of a Chronic Wound? JAMA 307 (2012).

8. S. E. Gardner, R. A. Frantz, B. N. Doebbeling, The validity of the clinical signs and symptoms used to identify localized chronic wound infection. Wound Repair and Regeneration: Official Publication of the Wound Healing Society [and] the European Tissue Repair Society 9, 178–186 (2001).

9. C. Giuliano, C. R. Patel, P. B. Kale-Pradhan, A Guide to Bacterial Culture Identification And Results Interpretation. Pharmacy and Therapeutics 44, 192 (2019).

10. N. D. Friedman, E. Temkin, Y. Carmeli, The negative impact of antibiotic resistance. Clinical Microbiology and Infection 22, 416–422 (2016).

11. B. A. Lipsky, et al., Antimicrobial stewardship in wound care: a Position Paper from the British Society for Antimicrobial Chemotherapy and European Wound Management Association. Journal of Antimicrobial Chemotherapy 71, 3026–3035 (2016).

12. T. Guinovart, G. Valdés-Ramírez, J. R. Windmiller, F. J. Andrade, J. Wang, Bandage-Based Wearable Potentiometric Sensor for Monitoring Wound pH. Electroanalysis 26, 1345–1353 (2014).

13. Y. Hattori, et al., Multifunctional Skin-Like Electronics for Quantitative, Clinical Monitoring of Cutaneous Wound Healing. Advanced Healthcare Materials 3, 1597–1607 (2014).

14. P. Mostafalu, et al., Smart Bandage for Monitoring and Treatment of Chronic Wounds. Small 14, 1703509 (2018).

15. S. RoyChoudhury, et al., Continuous Monitoring of Wound Healing Using a Wearable Enzymatic Uric Acid Biosensor. Journal of The Electrochemical Society 165, B3168–B3175 (2018).

16. E. Shirzaei Sani, et al., A stretchable wireless wearable bioelectronic system for multiplexed monitoring and combination treatment of infected chronic wounds. Science Advances 9 (2023).

17. H. Ryu, et al., Materials and Device Designs for Wireless Monitoring of Temperature and Thermal Transport Properties of Wound Beds during Healing. Advanced Healthcare Materials 13 (2023).

18. Y. Gao, et al., A flexible multiplexed immunosensor for point-of-care in situ wound monitoring. Science Advances 7 (2021).

19. M. Shirasu, K. Touhara, The scent of disease: volatile organic compounds of the human body related to disease and disorder. Journal of Biochemistry 150, 257–266 (2011).

20. M. Ashrafi, et al., Volatile organic compound detection as a potential means of diagnosing cutaneous wound infections. Wound Repair and Regeneration 25, 574–590 (2017).

21. A. N. Thomas, et al., Novel noninvasive identification of biomarkers by analytical profiling of chronic wounds using volatile organic compounds. Wound Repair and Regeneration 18, 391–400 (2010).

22. W. Filipiak, et al., Molecular analysis of volatile metabolites released specifically by staphylococcus aureus and pseudomonas aeruginosa. BMC Microbiology 12 (2012).

23. E. Daulton, A. Wicaksono, J. Bechar, J. A. Covington, J. Hardwicke, The Detection of Wound Infection by Ion Mobility Chemical Analysis. Biosensors 10, 19 (2020).

24. G. C. Gurtner, S. Werner, Y. Barrandon, M. T. Longaker, Wound repair and regeneration. Nature 453, 314–21 (2008).

25. D. Kundu, A. Jayaraman, C. K. Sen, Clinical Measurement of Transepidermal Water Loss. Advances in Wound Care 15 (2025).

26. D. Chattopadhyay, et al., Deficient functional wound closure as measured by elevated trans-epidermal water loss predicts chronic wound recurrence: An exploratory observational study. Scientific Reports 14 (2024).

27. J. Shin, et al., A non-contact wearable device for monitoring epidermal molecular flux. Nature 640, 375–383 (2025).

28. Y. Zhang, et al., Flexible integrated sensing platform for monitoring wound temperature and predicting infection. Microbial Biotechnology 14, 1566–1579 (2021).

29. D. J. Freeman, F. R. Falkiner, C. T. Keane, New method for detecting slime production by coagulase negative staphylococci. Journal of Clinical Pathology 42, 872–874 (1989).

30. P. Martin, Wound Healing--Aiming for Perfect Skin Regeneration. Science 276, 75–81 (1997).

31. A. J. Singer, R. A. F. Clark, Cutaneous Wound Healing. New England Journal of Medicine 341, 738–746 (1999).

32. E. Proksch, J. M. Brandner, J.-M. Jensen, The skin: an indispensable barrier. Experimental dermatology 17, 1063–72 (2008).

33. S. A. Eming, P. Martin, M. Tomic-Canic, Wound repair and regeneration: Mechanisms, signaling, and translation. Science Translational Medicine 6, 265sr6–265sr6 (2014).

34. I. Pastar, et al., Epithelialization in Wound Healing: A Comprehensive Review. Advances in Wound Care 3, 445–464 (2014).

35. N. X. Landén, D. Li, M. Ståhle, Transition from Inflammation to proliferation: a Critical Step during Wound Healing. Cellular and Molecular Life Sciences 73, 3861–3885 (2016).

36. T. Lucas, et al., Differential Roles of Macrophages in Diverse Phases of Skin Repair. The Journal of Immunology 184, 3964–3977 (2010).

37. M. G. Tonnesen, X. Feng, R. A. F. Clark, Angiogenesis in Wound Healing. Journal of Investigative Dermatology Symposium Proceedings 5, 40–46 (2000).

38. I. E. Johnson, T. A. Wilgus, Vascular Endothelial Growth Factor and Angiogenesis in the Regulation of Cutaneous Wound Repair. Advances in Wound Care 3, 647–661 (2014).

39. P. Krzyszczyk, R. Schloss, A. Palmer, F. Berthiaume, The Role of Macrophages in Acute and Chronic Wound Healing and Interventions to Promote Pro-wound Healing Phenotypes. Frontiers in Physiology 9 (2018).

40. D. M. Mosser, J. P. Edwards, Exploring the full spectrum of macrophage activation. Nature Reviews Immunology 8, 958–969 (2008).

41. I. D. J. Bos, P. J. Sterk, M. J. Schultz, Volatile Metabolites of Pathogens: A Systematic Review. PLoS Pathogens 9, e1003311 (2013).

42. S. Guo, L. A. DiPietro, Factors Affecting Wound Healing. Journal of Dental Research 89, 219–229 (2010).

43. H. Brem, M. Tomic-Canic, Cellular and molecular basis of wound healing in diabetes. Journal of Clinical Investigation 117, 1219–1222 (2007).

44. V. Falanga, Wound healing and its impairment in the diabetic foot. The Lancet 366, 1736–1743 (2005).

45. D. Clausen, et al., Wearable continuous diffusion-based skin gas analysis. Nature Communications 16 (2025).

46. J. Kim, A. S. Campbell, B. E.-F. de Ávila, J. Wang, Wearable biosensors for healthcare monitoring. Nature Biotechnology 37, 389–406 (2019).

47. J. Heikenfeld, et al., Accessing analytes in biofluids for peripheral biochemical monitoring. Nature Biotechnology 37, 407–419 (2019).

48. G. A. James, et al., Biofilms in chronic wounds. Wound Repair and Regeneration 16, 37–44 (2008).

49. I. Malone, et al., The Prevalence of Biofilms in Chronic wounds: a Systematic Review and meta-analysis of Published Data. Journal of Wound Care 26, 20–25 (2017).

50. W. Miekisch, J. K. Schubert, G. F. E. Noeldge-Schomburg, Diagnostic potential of breath analysis—focus on volatile organic compounds. Clinica Chimica Acta 347, 25–39 (2004).

51. A. W. Boots, et al., The versatile use of exhaled volatile organic compounds in human health and disease. Journal of Breath Research 6, 027108 (2012).

52. X. Huang, et al., A skin-interfaced three-dimensional closed-loop sensing and therapeutic electronic wound bandage. Nature Communications 16 (2025).

53. H. Li, et al., Towards adaptive bioelectronic wound therapy with integrated real-time diagnostics and machine learning–driven closed-loop control. npj Biomedical Innovations 2 (2025).

