## Supplementary material for "Healing cascades and infections in wounds monitored using a wearable sensor of gaseous flux": SI

**This PDF file includes:**

Supporting text
Figures S1 to S21
Tables S1 to S2
SI References

### Supporting Information Text

#### Modeling for Bacterial Growth and VOC Kinetics

**Bacterial growth kinetics: Geometric dependency**

Experimental observations indicate that bacterial growth kinetics are fundamentally governed by the initial seeding density, which dictates the spatial geometry of colony formation. To accurately capture these distinct physical regimes, two specific kinetic models are applied, reflecting the different nutrient availability and spatial constraints.

The high inoculation group is characterized by the rapid merger of closely spaced micro-colonies into a continuous, thin bacterial lawn. Due to the high surface-to-volume ratio of this thin film, individual bacteria maintain sufficient contact with the substrate, minimizing spatial diffusion gradients. This configuration approximates a quasi-homogeneous system similar to planktonic growth in liquid culture[1]. Consequently, this regime is modeled using the classical Monod equation extended with a decay term to account for the decrease in bacteria load[2,3].

$$\frac{dX}{dt} = \mu_{max} \frac{S}{K_s + S} X - k_d X \tag{1}$$

$$\frac{dS}{dt} = -\frac{1}{Y_{X/S}} \mu_{max} \frac{S}{K_s + S} X \tag{2}$$

Where $X$ represents bacteria load, $S$ is the substrate concentration, $\mu_{max}$ is the maximum specific growth rate, $K_s$ is the half-saturation constant, and $k_d$ is the specific death rate.

In contrast, the low inoculation group grows as discrete, large macro-colonies. Unlike the thin lawn in the high group, these large colonies suffer from significant spatial heterogeneity: nutrient depletion occurs rapidly in the dense center, restricting active growth to the outer periphery (the "active rim")[1,4]. Because the growth is geometrically limited to the edge, the system does not follow exponential Monod kinetics. Instead, following the colony kinetic theory proposed by Pirt (1967), it is assumed that the colony radius $r$ expands linearly with time after an initial adaptation period[4,5]. Consequently, the radial growth follows $r \propto (t - t_0)$, where $t_0$ represents the effective lag time. Assuming the colony maintains a relatively constant height, the total bacteria load $X$ scales with the colony area, yielding a quadratic kinetic model:

$$X(t) \propto r(t)^2 \propto (t - t_0)^2 \tag{3}$$

This model explicitly captures the "edge-growth" mechanism characteristic of solid-state limitations.

**VOC flux kinetics: Surface-saturation model**

Traditional Luedeking-Piret model[6] fails to describe the observed saturation of VOC flux at high bacteria load levels. To explain this non-linear behavior, a mechanistic model integrating Fickian diffusion and product inhibition is derived.

It is postulated that the specific VOC production rate $\beta$ is not constant but is subject to linear feedback inhibition by the local VOC concentration $P$ within the colony matrix, described as $\beta = \beta_0(1 - P/P_m)$, where $\beta_0$ is the intrinsic production coefficient and $P_m$ is the maximum product tolerance[6,7]. Simultaneously, the release of VOCs from the colony surface is governed by Fickian diffusion, modeled as $J_{VOC} = kP$, where $k$ represents the effective diffusion coefficient. Assuming the system operates under a quasi-steady state where the biological production rate balances the diffusive flux $\beta X = J_{VOC}$, the local product concentration can be expressed as a function of bacteria load:

$$P = \beta_0 X/(k + \beta_0 X/P_m) \tag{4}$$

Substituting this equilibrium concentration back into the flux equation yields a saturation-type kinetic function:

$$J_{VOC} = \frac{A \cdot X}{B + X} \tag{5}$$

where the lumped parameter $A = kP_m$ represents the maximum flux, and $B = kP_m/\beta_0$ defines the half-saturation bacteria load.

**Figures**

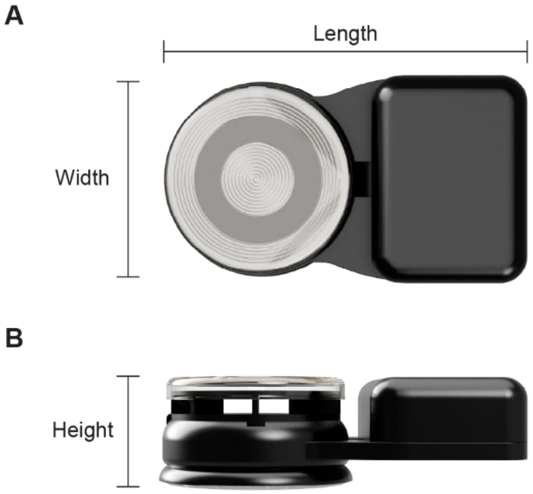


**Fig. S1.** Device dimensions. (*A*) Top-view and (*B*) side-view illustrations of the flux sensor measuring 22 mm in width, 40 mm in length, 14 mm in height, and 7.59 g in weight including a lithium polymer battery.

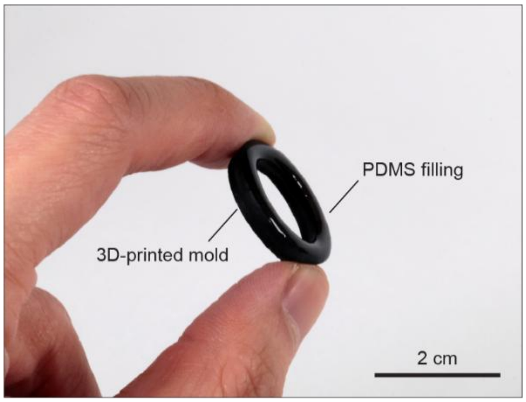


**Fig. S2.** Disposable adaptor for the flux sensor for wound monitoring.

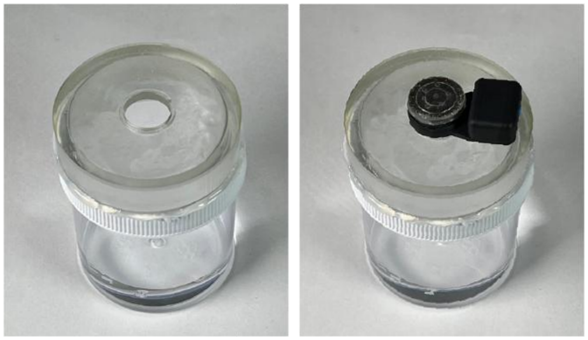

**Fig. S3.** Set-up for validation experiments showing a container filled with either water or an organic solvent and a lid with designs allowing the flux sensor to properly fit.

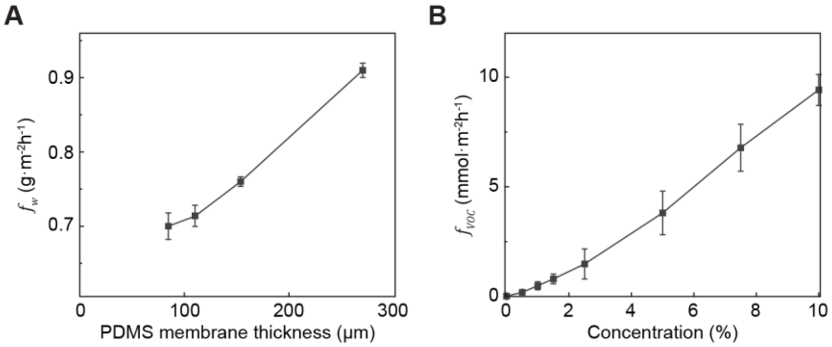


**Fig. S4.** Linearity of flux measurements. $\boldsymbol{f_w}$ (*A*) with various thicknesses of PDMS membranes as a diffusion barrier and $\boldsymbol{f_{VOC}}$ (*B*) during exposure to vapor from various concentrations of isopropyl alcohol. $\boldsymbol{f_w}$: Water vapor flux. $\boldsymbol{f_{VOC}}$: VOC flux.

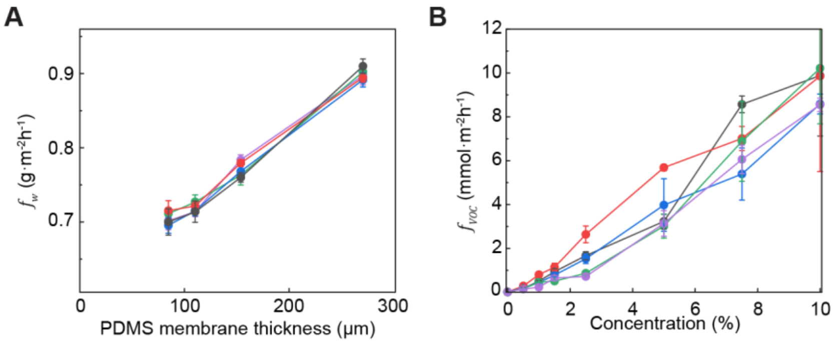


**Fig. S5.** Reproducibility of flux measurements. $\boldsymbol{f_w}$ (*A*) with various thicknesses of pdms membranes as a diffusion barrier and $\boldsymbol{f_{VOC}}$ (*B*) during exposure to vapor from various concentrations of isopropyl alcohol across five different days. $\boldsymbol{f_w}$: Water vapor flux. $\boldsymbol{f_{VOC}}$: VOC flux.

**A** $10^3$ CFU

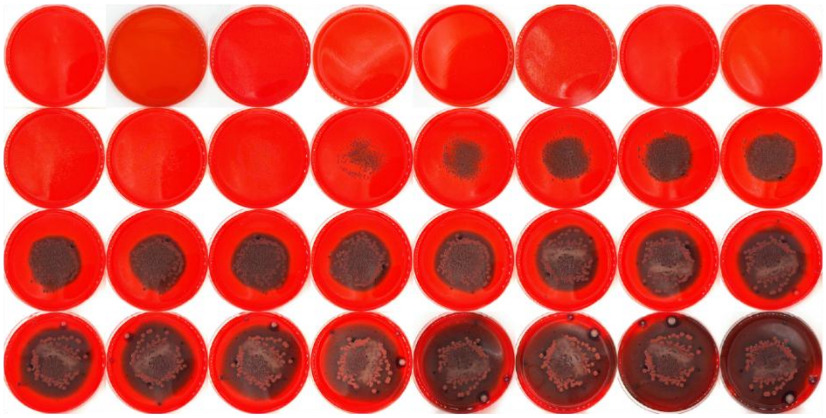

**B** $10^7$ CFU

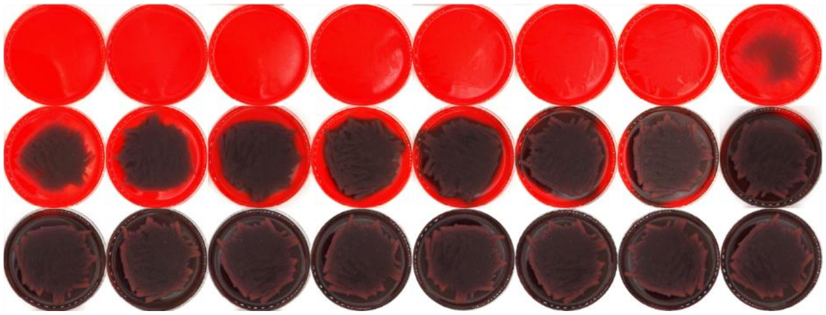

**Fig. S6.** Images of biofilm culture plates inoculated with $10^3$ CFU (*A*) and $10^7$ CFU (*B*) *S. aureus* taken every 6 h from 0 to 198 h.

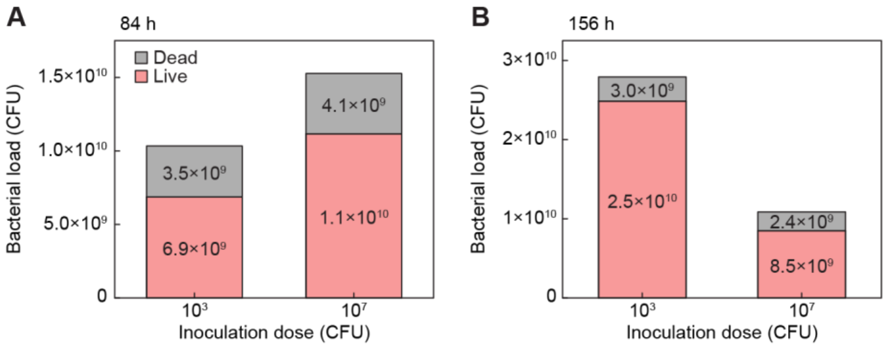


**Fig. S7.** Live/Dead stain results from biofilm culture plates at 84 h (*A*) and 156 h (*B*).

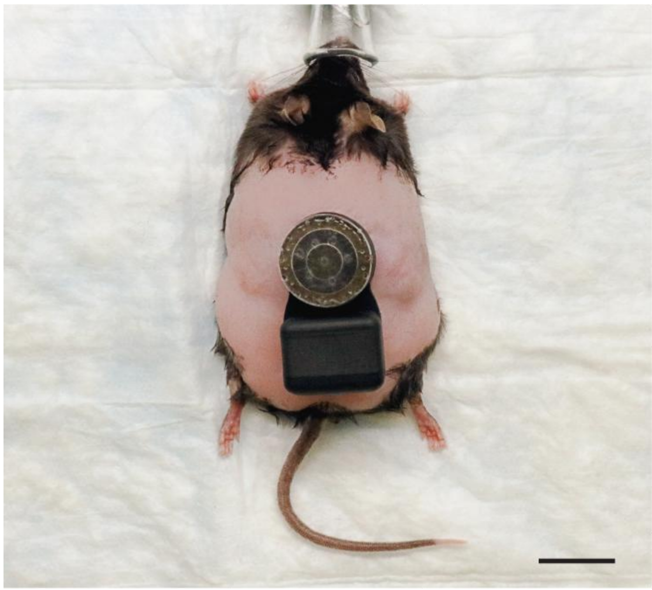

**Fig. S8.** Image of diabetic mouse with sensor placed on wound. Scale bar, 2 cm.

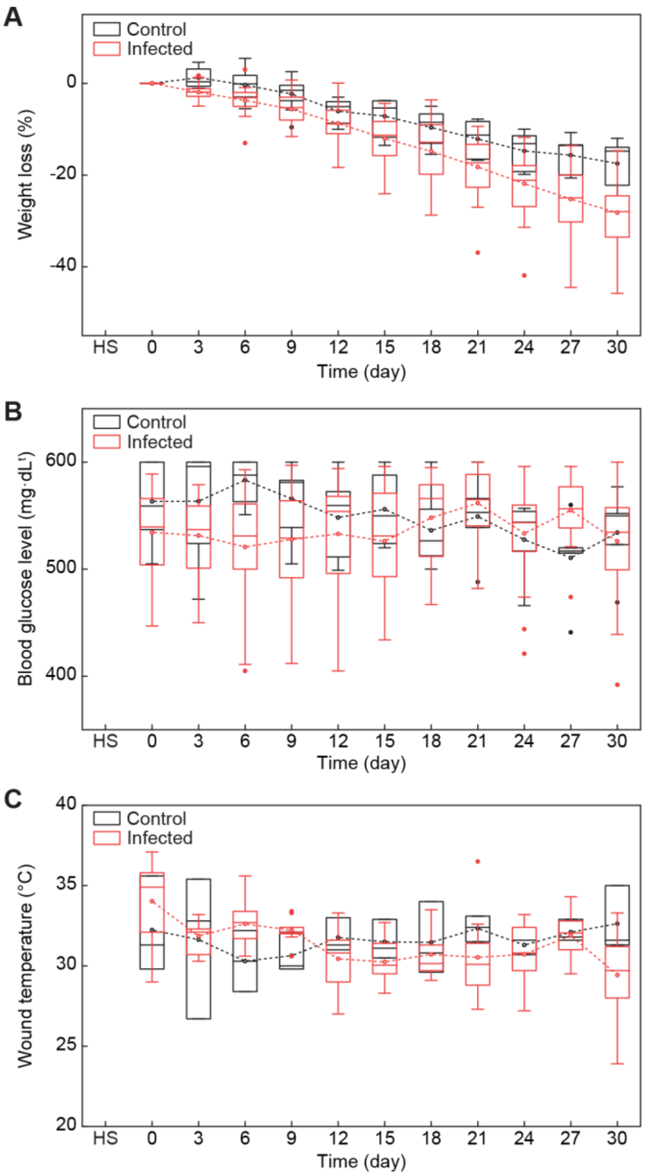


**Fig. S9.** Biological characterization of mice. For weight loss (*A*) and blood glucose (*B*), $n$ = 11 for days 0-9, $n$ = 8 for days 12-18, $n$ = 5 for days 21-30 for control and $n$ = 30 for days 0-9, $n$ = 27 for days 12-18, $n$ = 24 for days 21-30 for infected. For wound temperature (*C*), control $n$ = 3, infected $n$ = 10. Horizontal line within the box represents the sample median and the ends of the box correspond to the interquartile range (first to third quartiles). The whiskers extend beyond the ends of the box by 1.5× (Interquartile range). Open symbols within the box represent the average and filled symbols outside the whiskers indicate outliers.

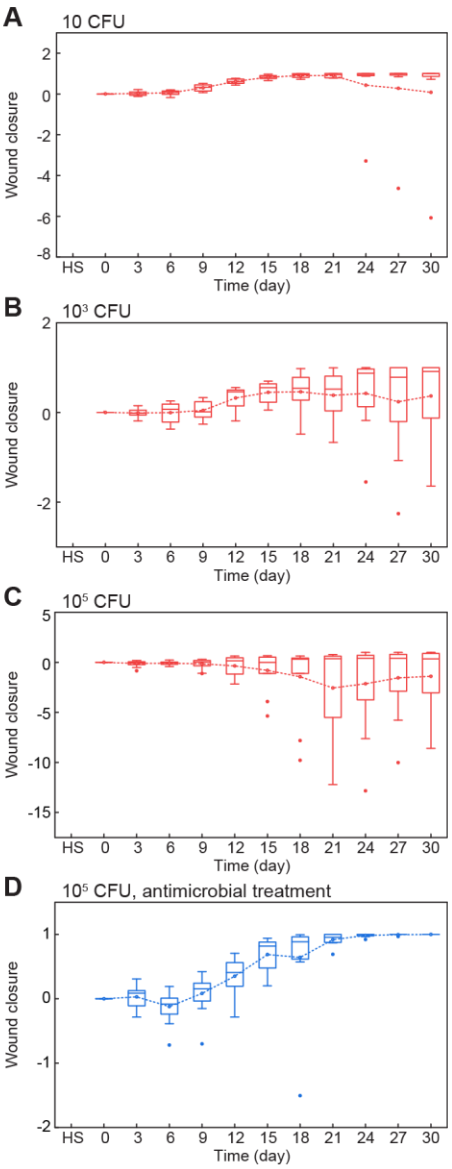


**Fig. S10.** Wound closure from infected groups with different inoculation doses of 10 (*A*), $10^3$ CFU (*B*), $10^5$ CFU (*C*), and $10^5$ CFU with antimicrobial treatment (*D*). HS: healthy skin.

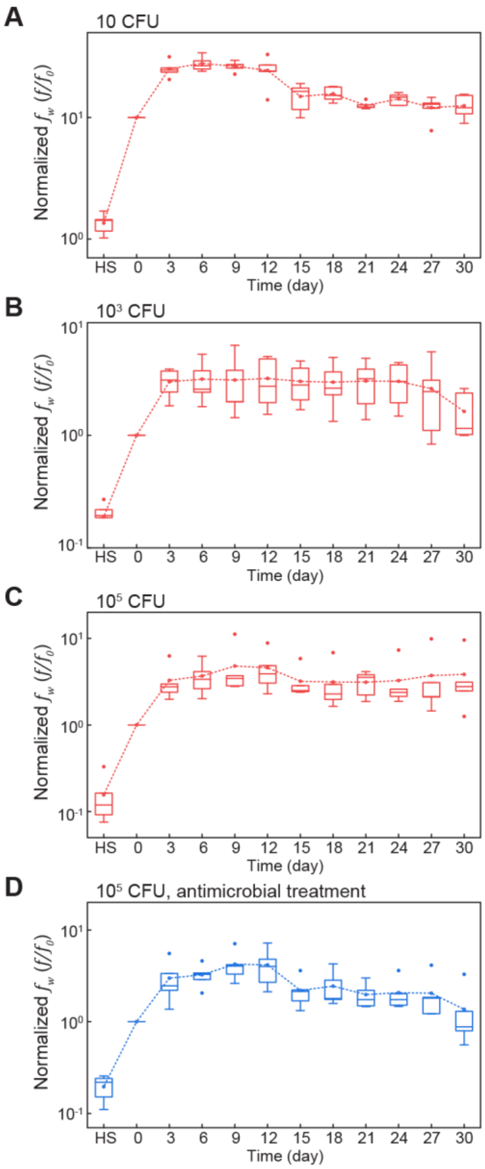


**Fig. S11.** Normalized $\boldsymbol{f_w}$ from infected groups with different inoculation doses of 10 (*A*), $10^3$ CFU (*B*), $10^5$ CFU (*C*), and $10^5$ CFU with antimicrobial treatment (*D*). $\boldsymbol{f_w}$: Water vapor flux. HS: Healthy skin.

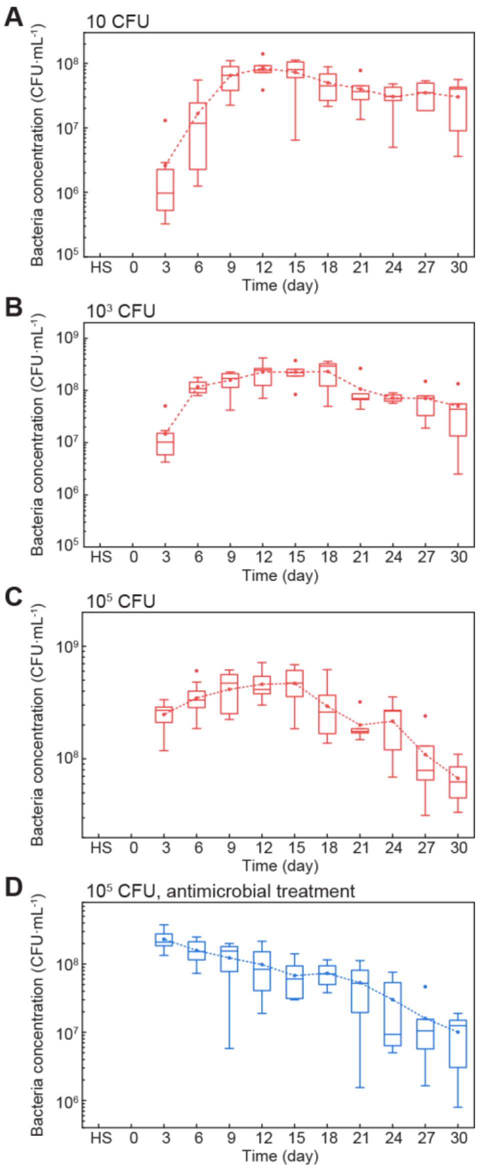


**Fig. S12.** Bacteria concentration from infected groups with different inoculation doses of 10 (*A*), $10^3$ CFU (*B*), $10^5$ CFU (*C*), and $10^5$ CFU with antimicrobial treatment (*D*). HS: healthy skin.

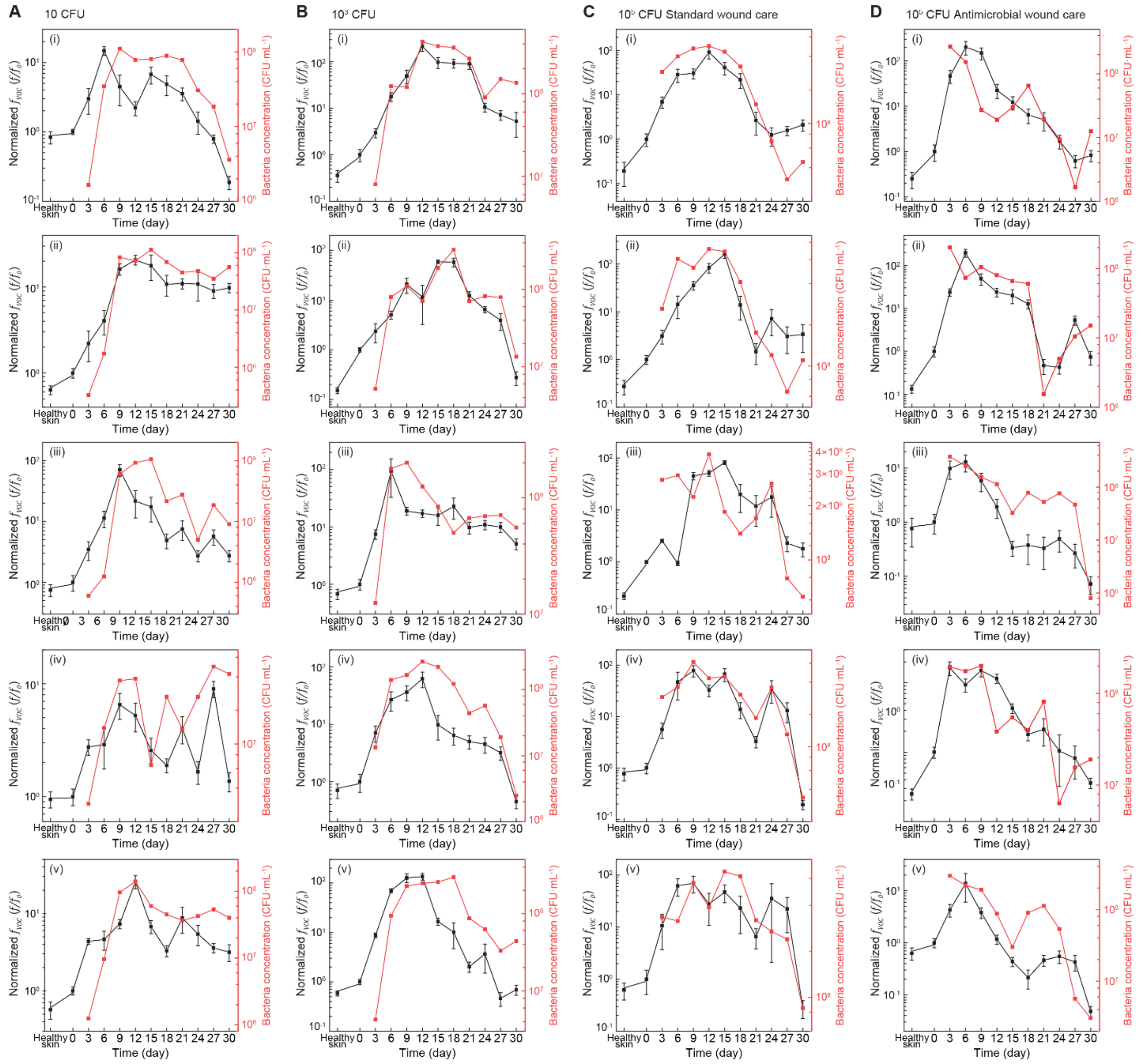


**Fig. S13.** Normalized $f_{VOC}$ compared with bacteria concentrations for individual samples from different infected groups with inoculation doses of 10 CFU (*A*), $10^3$ CFU (*B*), $10^5$ CFU with standard wound care (*C*), and $10^5$ CFU with antimicrobial treatment (*D*). $f_{VOC}$: VOC flux.

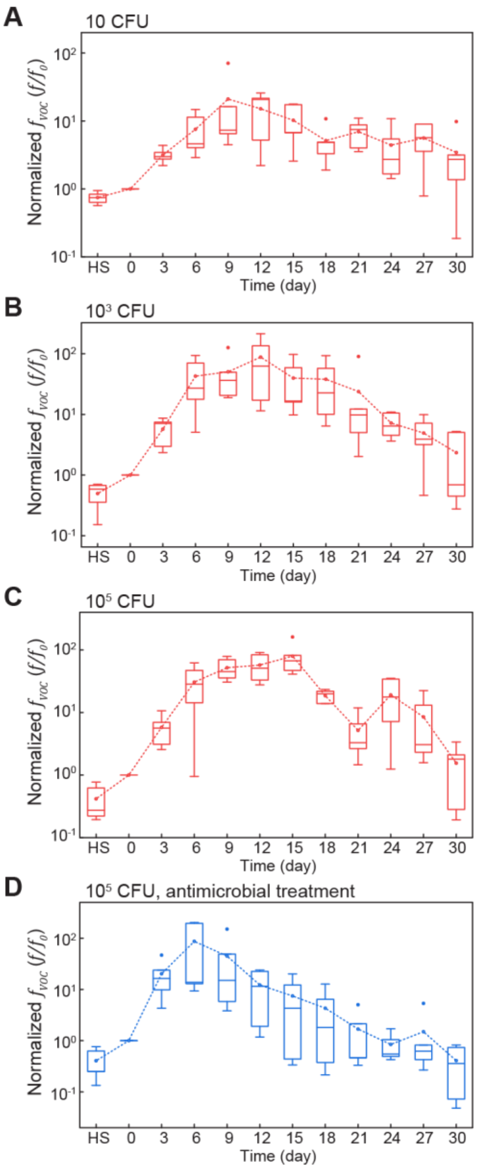


**Fig. S14.** Normalized $\boldsymbol{f_{VOC}}$ from infected groups with different inoculation doses of 10 CFU (*A*), $10^3$ CFU (*B*), $10^5$ CFU (*C*), and $10^5$ CFU with antimicrobial treatment (*D*). $\boldsymbol{f_{VOC}}$: VOC flux.

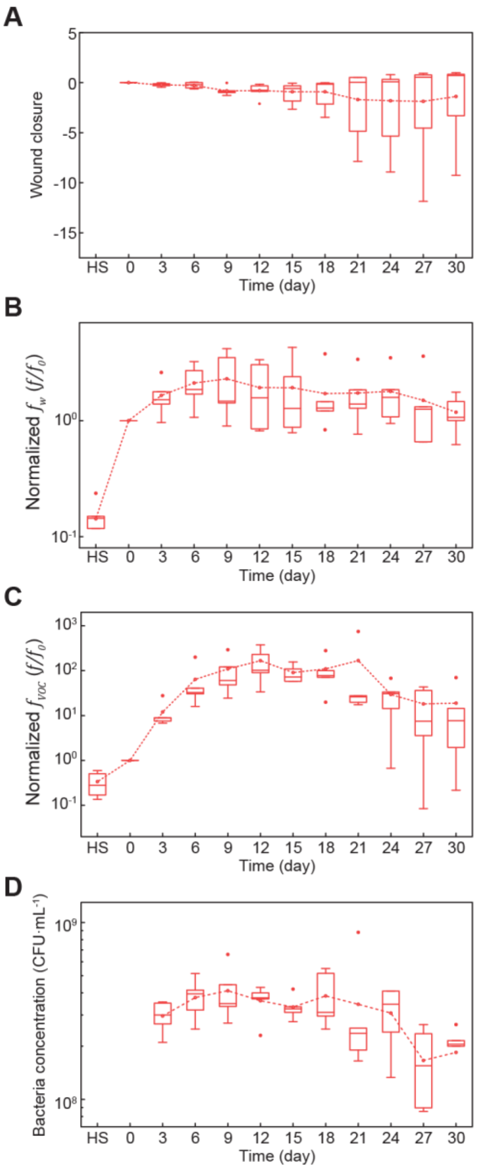


**Fig. S15.** Wound closure (*A*), normalized $\boldsymbol{f_w}$ (*B*), normalized $\boldsymbol{f_{VOC}}$ (*C*), and bacteria concentration (*D*) from an infected group with an inoculation dose of $10^7$ CFU. $\boldsymbol{f_w}$: Water vapor flux. $\boldsymbol{f_{VOC}}$: VOC flux. HS: Healthy skin.

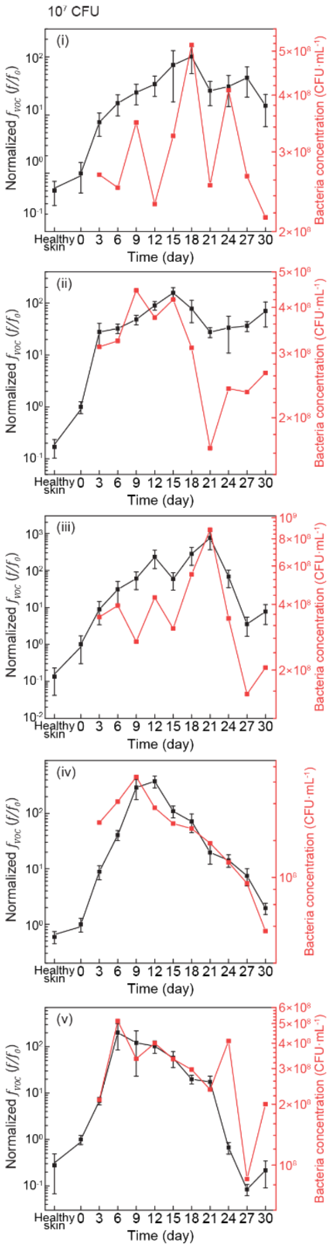


**Fig. S16.** Normalized $f_{VOC}$ compared with bacteria concentrations for individual samples from different infected groups with inoculation doses of $10^7$ CFU. $f_{VOC}$: VOC flux.

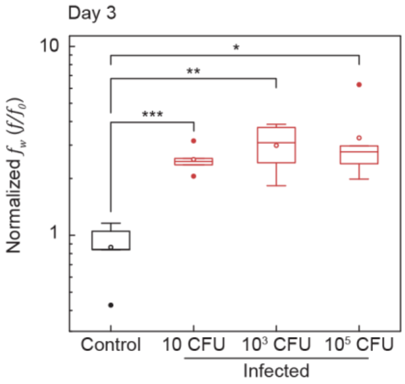


**Fig. S17.** Normalized $\boldsymbol{f_w}$ from the wound on day 3 for control (black) and infected (red) groups. *n* = 5 per group. ***, **, and * indicate $p < 0.001$, $p < 0.01$, and $p < 0.05$, respectively. $\boldsymbol{f_w}$: Water vapor flux.

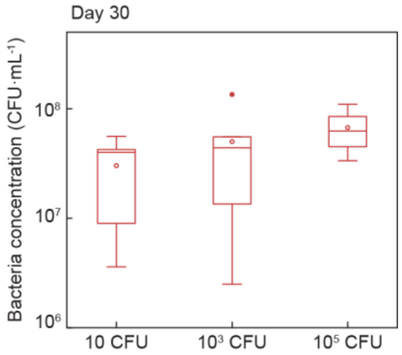


**Fig. S18.** Bacteria concentration in wounds of mice with inoculation doses of 10, $10^3$ CFU, and $10^5$ CFU on day 30. $n$ = 5 per group.

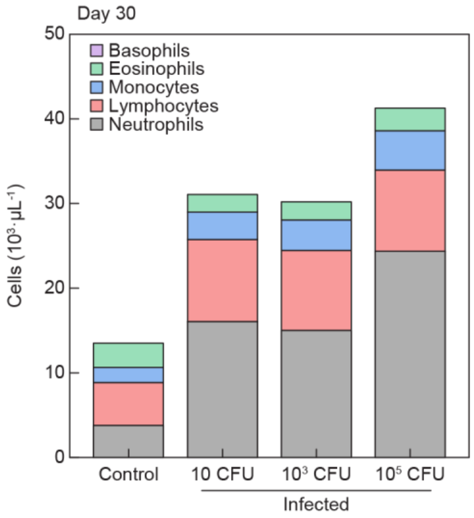


**Fig. S19.** White blood cell count and concentration of subtypes on day 30 compared between control and infected groups with various inoculation doses. Subtypes include neutrophils, lymphocytes, monocytes, eosinophils, and basophils. *n* = 5 control, *n* = 8 per infected group.

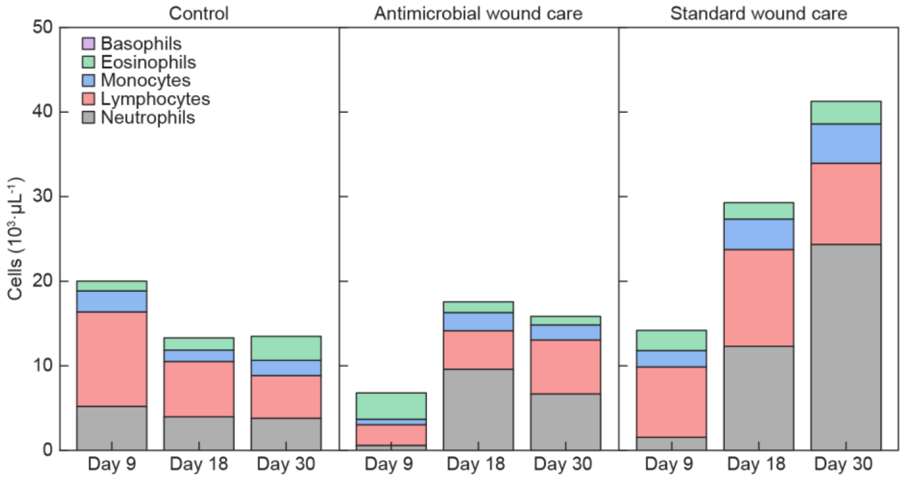


**Fig. S20.** White blood cell count and concentration of subtypes of control mice and infected groups with inoculation doses of $10^5$ CFU with antimicrobial wound care and standard wound care on days 9, 18, and 30. Subtypes include neutrophils, lymphocytes, monocytes, eosinophils, and basophils. *n* = 3 per group for days 9 and 18, *n* = 5 control and *n* = 8 per infected group day 30.

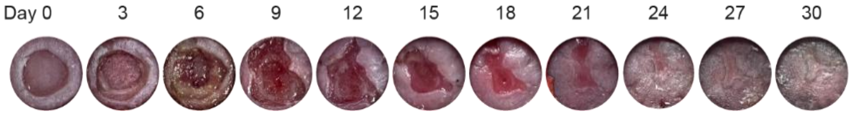


**Fig. S21.** Representative images of infected wounds of $10^5$ CFU inoculation with antimicrobial wound care recorded every 3 days.

**Tables**

**Table S1.** Inoculation dose of in vitro samples

| Sample ID | Inoculation (CFU) |
|---|---|
| CR-3 | $1.95 \times 10^3$ |
| CR-7 | $1.827 \times 10^7$ |

**Table S2.** Inoculation dose of individual mice

| Mouse ID | Inoculation (CFU) |
|---|---|
| CONTROL-1 | - |
| CONTROL -2 | - |
| CONTROL -3 | - |
| CONTROL -4 | - |
| CONTROL -5 | - |
| 1-SWC-1 | 1.95 × 10 |
| 1-SWC-2 | 1.95 × 10 |
| 1-SWC-3 | 1.98 × 10 |
| 1-SWC-4 | 1.98 × 10 |
| 1-SWC-5 | 1.98 × 10 |
| 3-SWC-1 | $2.1 \times 10^3$ |
| 3-SWC-2 | $1.53 \times 10^3$ |
| 3-SWC-3 | $1.98 \times 10^3$ |
| 3-SWC-4 | $1.98 \times 10^3$ |
| 3-SWC-5 | $1.98 \times 10^3$ |
| 5-SWC-1 | $1.65 \times 10^5$ |
| 5-SWC-2 | $1.65 \times 10^5$ |
| 5-SWC-3 | $1.95 \times 10^5$ |
| 5-SWC-4 | $1.95 \times 10^5$ |
| 5-SWC-5 | $1.95 \times 10^5$ |
| 5-AWC-1 | $1.65 \times 10^5$ |
| 5-AWC-2 | $1.65 \times 10^5$ |
| 5-AWC-3 | $1.95 \times 10^5$ |
| 5-AWC-4 | $1.95 \times 10^5$ |
| 5-AWC-5 | $1.95 \times 10^5$ |
| 7-SWC-1 | $1.785 \times 10^7$ |
| 7-SWC-2 | $1.785 \times 10^7$ |
| 7-SWC-3 | $1.815 \times 10^7$ |
| 7-SWC-4 | $1.815 \times 10^7$ |
| 7-SWC-5 | $1.761 \times 10^7$ |

**SI References**


1. Ranade, S., Zhang, Y., Kaplan, M., Majeed, W. & He, Q. Metabolic Engineering and Comparative Performance Studies of Synechocystis sp. PCC 6803 Strains for Effective Utilization of Xylose. *Frontiers in Microbiology* **6**, (2015).
2. Monod, J. The Growth of Bacterial Cultures. *Annual Review of Microbiology* **3**, 371–394 (1949).
3. Fuad, M. N. M. Application of modified Logistic and Monod models in a single equations system framework to improve cell culture growth modeling and estimation. *Authorea* (2020) doi:https://doi.org/10.22541/au.160647305.55974012/v3.
4. PIRT, S. J. A Kinetic Study of the Mode of Growth of Surface Colonies of Bacteria and Fungi. *Journal of General Microbiology* **47**, 181–197 (1967).
5. Cooper, A. L., Dean, A. C. R. & Hinshelwood, C. N. Factors affecting the growth of bacterial colonies on agar plates. *Proceedings of the Royal Society of London. Series B. Biological Sciences* **171**, 175–199 (1968).
6. Luedeking, R. & Piret, E. L. A kinetic study of the lactic acid fermentation. Batch process at controlled pH. *Biotechnology and Bioengineering* **67**, 636–644 (2000).
7. Levenspiel, O. The monod equation: A revisit and a generalization to product inhibition situations. *Biotechnology and Bioengineering* **22**, 1671–1687 (1980).
